# Machine learning can be as good as maximum likelihood when reconstructing phylogenetic trees and determining the best evolutionary model on four taxon alignments

**DOI:** 10.1101/2023.07.12.548770

**Authors:** Nikita Kulikov, Fatemeh Derakhshandeh, Christoph Mayer

## Abstract

Phylogenetic tree reconstruction with molecular data is important in many fields of life science research. The gold standard in this discipline is the phylogenetic tree reconstruction based on the Maximum Likelihood method. In this study, we explored the utility of neural networks to predict the correct model of sequence evolution and the correct topology for four sequence alignments. We trained neural networks with different architectures using simulated nucleotide and amino acid sequence alignments for a wide range of evolutionary models, model parameters and branch lengths. By comparing the accuracy of model and topology prediction of the trained neural networks with Maximum Likelihood and Neighbour Joining methods, we show that for quartet trees, the neural network classifier outperforms the Neighbour Joining method and is in most cases as good as the Maximum Likelihood method to infer the best model of sequence evolution and the best tree topology. These results are consistent for nucleotide and amino acid sequence data. Furthermore, we found that neural network classifiers are much faster than the IQ-Tree implementation of the Maximum Likelihood method. Our results show that neural networks could become a true competitor for the Maximum Likelihood method in phylogenetic reconstructions.

## Introduction

In parallel with the development of sequencing technologies, many algorithms have been developed to reconstruct phylogenetic trees from molecular sequence data (Yang, 2014). Tree reconstruction methods fall into two main categories: distance-based methods such as UPGMA (Sneath & Sokal, 1973), Neighbour Joining (NJ) (Saitou & Nei, 1987) or minimum evolution (ME) (Edwards & Cavalli-Sforza, 1964), and character-based methods like Maximum Parsimony (MP) (Farris, 1970; Fitch, 1971), Maximum Likelihood (ML) (Cavalli-Sforza & Edwards, 1967; Felsenstein, 1981) and Bayesian inference methods (Ronquist & Huelsenbeck, 2003). These methods differ not only in methodology and underlying assumptions but also in crucial aspects such as statistical consistency, efficiency, and robustness (Penny et al., 1992). The capacity of a statistical method to accurately infer the correct phylogenetic tree, when the amount of data approaches infinity, is known as statistical consistency. A method is called efficient if the amount of data necessary to find the correct solution with a high probability is small and it is called robust if it finds the correct solution even if some of its underlying assumptions are slightly violated.

Today the most widely used method in phylogenetic reconstructions is the ML method. This method is under realistic conditions superior to distance-based and the MP method (Xuhua, 2018). The likelihood of a data set is the probability to observe this data by chance for a given model of sequence evolution, model parameters and a given tree. The tree and model that have the highest likelihood of producing the data set should be preferred and the aim of the ML methods is to find this model and tree. The need to assume a model of sequence evolution is occasionally criticised (Abadi et al., 2019), although they are inherently needed for all probability-based statistical methods. The ML approach has been proven to be statistically consistent (Graur & Li, 1997; Truszkowski & Goldman, 2016) under the premise that the evolutionary process (i.e. the model of sequence evolution) that created the data is among the evolutionary processes considered in the phylogenetic tree reconstruction. Beyond consistency, it has been shown that the ML method is efficient as well as robust for a range of possible model violations (Yang, 2014).

Models of sequence evolution specify the probability that nucleotides or amino acids are substituted by other residues in given intervals of time. A wide range of mostly time reversible models of sequence evolution are used today (Felsenstein, 2004; Yang, 2014). For nucleotides, time reversible models range from the simplest possible model, the Jukes Cantor (JC) model (Jukes & Cantor, 1969) which assumes equal base frequencies and substitution rates for all nucleotides, to the most general time reversible (GTR) model (Tavaré, 1986). For amino acids, usually models are used with empirical amino acid frequencies and substitution rates since estimating all parameters of the most general time reversible model with 20 residues is very time consuming (Whelan & Goldman, 2001; Keane et al., 2006). During a tree reconstruction, the model and its parameters are typically estimated along with the tree. Alternatively, if they are at least partially known they can be specified in advance.

There is theoretical evidence that no method can outcompete the ML method when the amount of available data approaches infinity. This follows from the Cramer-Rao lower bound theorem (Rao, 1945; Cramer, 1946) which provides an asymptotic lower bound for the achievable variance of consistent and unbiased estimators as the amount of data approaches infinity, together with the observation that the consistent and unbiased maximum likelihood estimator takes on this lower bound under relatively mild conditions asymptotically (Stuart et al. 1999, Chapter 18; Yang, 2014). In particular this means that no other estimator exists that is more efficient, i.e. requires less data, than the maximum likelihood estimator used in phylogenetic reconstructions. Altogether our goal cannot be to find a method that has statistical properties that are better than those of the ML method, but a method that is equally good and computationally more efficient.

The traditional tree reconstruction methods have different advantages and disadvantages. Distance based methods normally require fewer computational resources but are less efficient and therefore less accurate than the ML method under realistic conditions (Huelsenbeck, 1995), while the ML method is computationally intensive but efficient and statistically consistent.

In recent years machine learning has become an important method not only in many fields of data science in general but also in biology (Borowiec et al., 2022). Machine learning refers to a wide range of methods that can be used mainly to classify data or solve regression problems. Apart from the Neural Networks (NNs), this includes popular methods such as decision trees, Random Forest (RF), Support Vector Machine (SVM), and k-means clustering (Mahesh, 2020). As machine learning methods are particularly good at solving classification problems, they should be candidates to classify/predict models of sequence evolution and topologies for given alignments of molecular sequence data. Indeed, machine learning has been proposed for phylogenetic tree reconstruction as early as in 2012 (Halgaswaththa et al., 2012). Several studies concentrated on inferring the correct topology of quartet trees by analysing the image created from aligned sequences, i.e. after representing the nucleotides with different colours and applying image recognition techniques to these images using Convolutional Neural Networks (CNNs, LeCun et al., 1989) with a large number of convolutional layers (Suvorov et al., 2020; Zou et al., 2020; Suvorov & Schrider, preprint 2022). Image classification of alignments using deep CNNs has also been suggested for determining the best evolutionary model of sequence evolution (Burgstaller-Muehlbacher et al., preprint 2021). Furthermore, Pinheiro et al. (2022) used a RF algorithm for predicting missing sequence regions and using this information to reconstruct phylogenetic trees. However, the accuracy of the RF method could hardly come close to that of the Neighbour Joining method.

Machine learning has also been proposed for phylogenetic tree reconstruction in combination with alignment free methods. Zhu & Cai (2021) used k-mer frequencies to represent biological sequences and used them to train a reinforcement learning model. Unfortunately, they compared their results with distance-based methods such as the UPGMA method rather than the ML method. Similarly, Gamage et al. (2020) used k-mer frequencies as a proxy for phylogenetic distances in combination with RF algorithm to infer the tree topology. The results were compared to the NJ method.

Several machine learning algorithms proposed so far are inferior to the ML method and only as good as distance-based methods (Zhu & Cai, 2021; Pinheiro et al., 2022). Suvorov et al. (2020) and Zou et al. (2020) showed that an image-based alignment classification with CNNs can yield good results. Burgstaller-Muehlbacher et al. (preprint 2021) have shown that CNNs analysing images of nucleotide alignments can compete with the ML method when it comes to model prediction and computing times.

Here we propose supervised machine learning using NNs as an alternative to existing model selection and topology reconstruction methods. The NNs we propose use site pattern frequencies (see Fig. 1) of four sequence alignments to predict the best model of sequence evolution and the most likely tree topology that created the alignment. Site pattern frequencies contain the full information of an alignment, if alignment sites are treated as independent and identically distributed as assumed in the ML method for all commonly used models of sequence evolution. Instead of computing the likelihood function for the set of alignment sites, we trained the NNs such that they can predict the best model and topology from the site pattern frequency distributions. If sufficiently complex NNs are trained with data sets that sample the full variety of possible data sufficiently dense, there is no reason this method could not achieve the accuracy of the ML method. We expect higher reconstruction accuracies when topology prediction networks were specifically trained for the different nucleotide and amino acid evolutionary models. Therefore, we trained topology prediction networks separately for a set of evolutionary models of sequence evolution. Analogously to the ML method, this implies that the best model is estimated prior to the final topology inference. An interesting property of NNs based on dense layers is that after the pattern frequencies have been determined, a prediction requires constant computational resources. Considering the computational intensity required to evaluate the likelihood function even for a small number of taxa, the number of operations needed to determine the network response for a single input should be relatively low. Therefore NNs should be computationally more efficient than the ML method for model and topology predictions using site pattern frequencies.

**Fig. 1.**
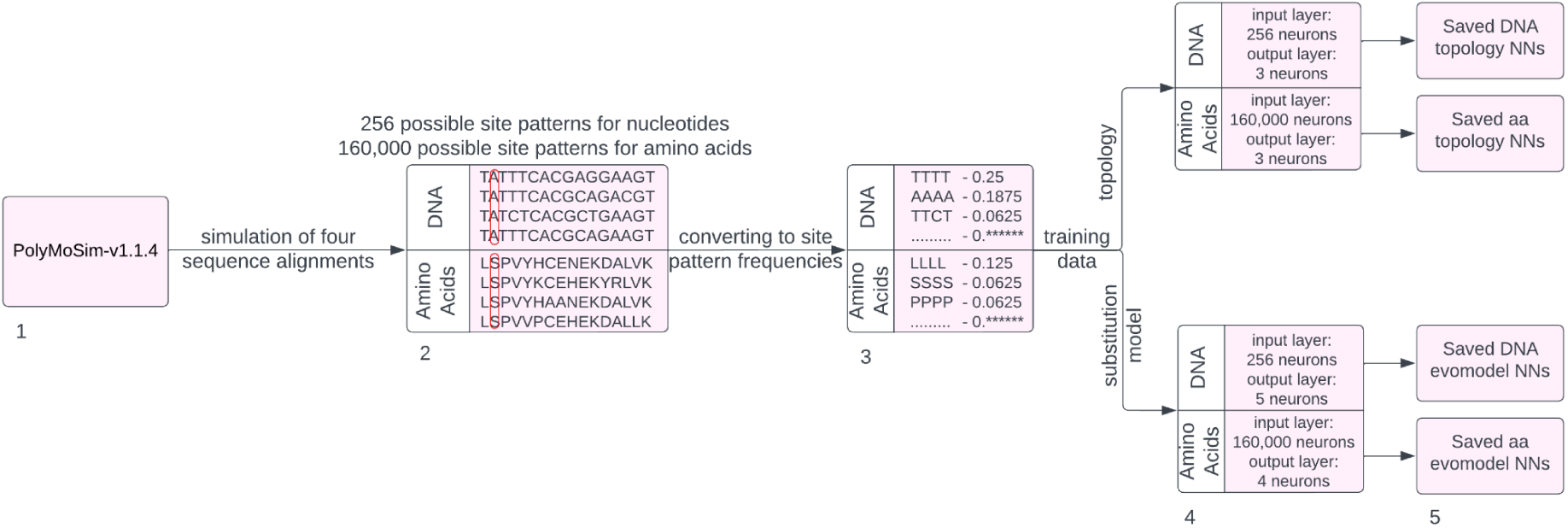
Schematic flowchart for simulating data sets and training NNs. (1) Alignment simulation. (2) Storing correct classification and site pattern frequencies. (3-4) Training of NNs with known classification and site pattern frequencies data. (5) Saving trained NNs. Note that site patterns are defined as the ordered strings of all nucleotide or amino acid residues at given alignment positions. Site pattern frequencies are the relative frequencies with which site pattern strings occur in the alignment. In the shown nucleotide alignment the site pattern “TTCT” occurs at alignment position 4 and has a frequency of 1/16.

In parallel and independent of us, Suvorov & Schrider (preprint 2022) have developed NNs that use the site pattern frequency distribution of four nucleotide sequences for training NNs to predict branch lengths. They used images of four taxon alignments for topology classifications and site pattern frequencies of four taxon alignments for branch length estimates and found that these estimates are about as good as those of the ML method. Leuchtenberger et al. (2020) proposed site pattern frequencies in combination with NNs to select the best tree reconstruction method for the Farris and Felsenstein Zone.

A limitation of using machine learning for phylogenetic reconstruction, whether we analyse site pattern frequencies or images of alignments, is that each possible classification has to be trained with sufficient training data, i.e. for all topologies we need to consider all combinations of branch lengths, models and model parameters. Therefore, we restrict our analyses to the four taxon case in which only three topologies are possible (Huelsenbeck & Hillis, 1993).

Here, we describe a set of six NN architectures and how they can be trained to predict the best model of sequence evolution and the best tree topology for four taxon alignments of nucleotide or amino acid sequences. Since we use site pattern frequencies to classify models and topologies, the input layers of the NNs have a number of neurons equal to the number of different site patterns in four taxon alignments, i.e. 256=4^4^ for nucleotides and 160,000=20^4^ for amino acids (Table 1). The NNs are trained by presenting them a large number of pattern frequency vectors together with the correct classification, i.e. the correct model of sequence evolution or the correct topology under which the four taxon alignment was created. Finally, the number of neurons in the output layer has to be equal to the number of different classes in the prediction problem. Specifically, the output values of these neurons are equal to the probabilities the NN assigns to the different possible outcomes. For the training we use simulated alignments that evolved under known hypotheses, since only for these the true topology and model are certainly known and only for these a sufficient number of training data sets is available that sample the full variety of each prediction class of our classification problem.

**Table 1:** Sizes of input and output layers and information on NN training.

|  | **Nucleotide**<br>**model**<br>**prediction** | **Nucleotide**<br>**topology**<br>**prediction** | **Amino acid**<br>**model**<br>**prediction** | **Amino acid**<br>**topology**<br>**prediction** |
| --- | --- | --- | --- | --- |
| **Neurons in input**<br>**layer** | 256 | 256 | 160,000 | 160,000 |
| **Neurons in**<br>**output layer, i.e.**<br>**number of**<br>**classes in**<br>**prediction** | 5 | 3 | 4 | 3 |
| **Length of**<br>**training**<br>**alignments** | 30,000 bp | 100,000 bp | 1,000,000 aa | 1,000,000 aa |
| **Alignments with**<br>**normal internal**<br>**branch length** | 450,000 | 300,000 | 150,000 | 30,000 |
| **Alignments with**<br>**short internal**<br>**branch length** | 450,000 | 300,000 | 150,000 | 30,000 |
| **Models included** | JC, K80, F81,<br>HKY, GTR | JC, K80, F81,<br>F84, HKY, GTR | JTT, LG, WAG,<br>Dayhoff | JTT, LG, WAG,<br>Dayhoff |
| **Size of training**<br>**data set array** | (900,000; 256) | (600,000; 256) | (300,000;<br>160,000) | (60,000; 160,000) |
| **Number of NN**<br>**architectures** | 6 | 6 | 6 | 6 |
| **Total number of trained NNs** | 6 architectures | 6 architectures<br>* 6 models | 6 architectures | 6 architectures<br>* 4 models |

We also tested non-neural network classifiers such as nearest neighbours, SVM or RF for predicting the best tree topology. These classifiers are commonly used to classify frequency distributions (see e.g. Hastie et al., 2009) and could be a potential alternative to NN classifiers for our prediction tasks.

Finally, we compared the prediction success of the NNs we proposed with the success of the ML+BIC (Bayesian Information Criterion; Schwarz, 1978) method for predicting the best model of sequence evolution as well as that of the ML and Neighbour Joining method for predicting the best tree topology.

## Methods

In this study, we trained NNs to predict the model of sequence evolution and the unrooted tree topology that most likely created a given four-taxon alignment of nucleotide or amino acid sequences.

### Neural network architectures and implementation

The training and prediction scripts and the NNs were implemented in Python using the TensorFlow library (Abadi et al., 2016). Workflows for training the NNs and for using trained NNs for predictions are illustrated in Figs. 1-2. Distributions, such as site pattern frequencies, are best classified with a series of fully connected layers, called dense layers in TensorFlow. We designed six different NN architectures, which all consist of dense, normalisation and dropout layers, but with different numbers of branches leading from the input to the output layer and one network with interconnections. The NNs we implemented are described in detail in the supplementary materials Section 1 and in Figs. S1-S2. We refer to the six NN architectures as B1, B2, B3, B10, U, CU, where the n in Bn, stands for the number of independent branches in the network that lead from the input to the output layer, U stands for a U-shaped network and CU for a U-shaped network with internal connections. The U and CU models were introduced with the intention to test networks with a large number of free parameters in a non-trivial architecture in the hope that they are able to store more information about our complex frequency distributions. With its interconnections, the CU model shares some distant similarity with the U-Net proposed by Ronneberger et al., (2015) for image segmentation. However descending and ascending branches serve very different tasks here and it was merely introduced since we found that this architecture trains very fast and efficiently compared to the complexity and the large number of free parameters it has. The architecture is explained in more detail in the supplementary materials, sections 1.3-1.4. We tested the same six NN architectures for all classification tasks so that for each task the architecture with the most appropriate complexity can be chosen.

**Fig. 2.**
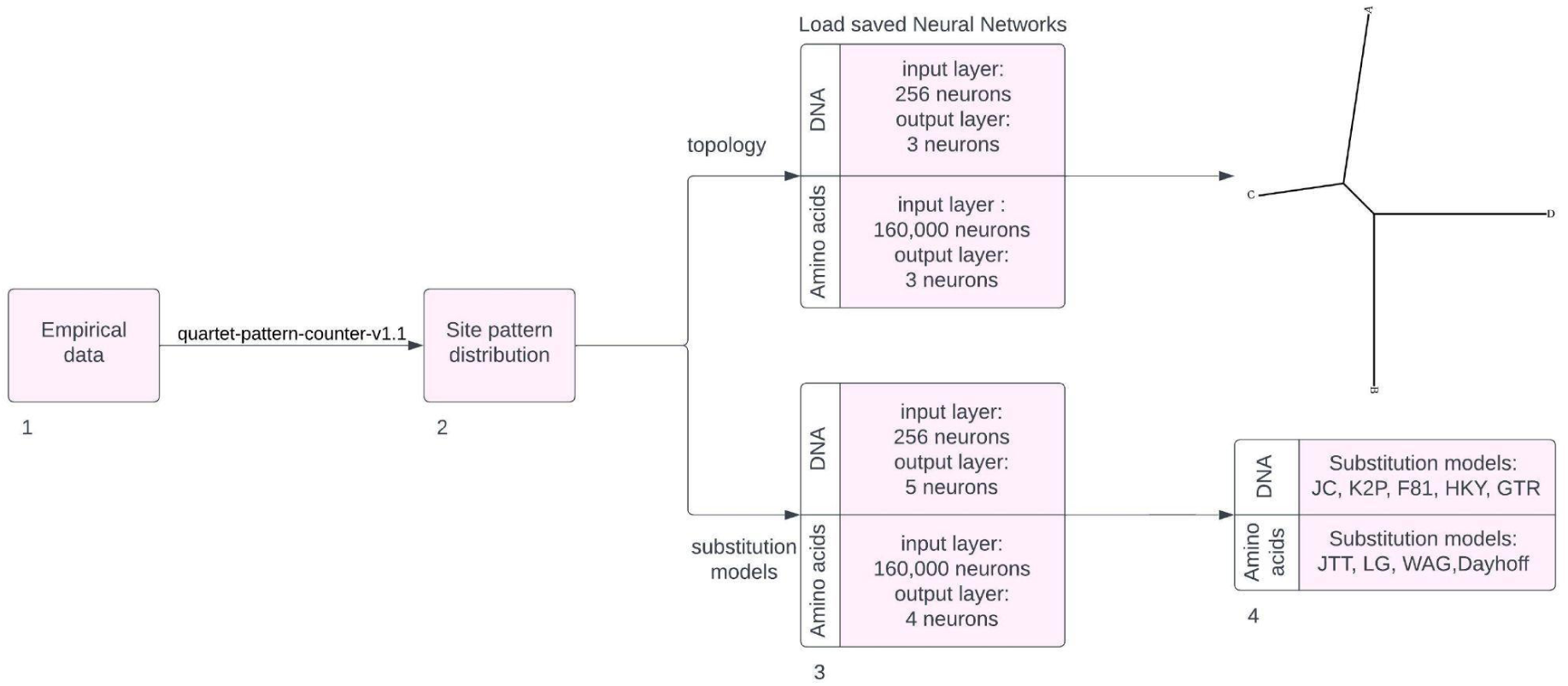
Schematic flowchart for model and topology predictions of empirical datasets using trained and saved NNs. (1) Loading alignment which shall be classified. (2) Determine pattern frequencies with Quartet-pattern-counter-v1.1 program. (3) Load trained NNs. (4) Classify alignment according to classification task. Model selection with subsequent topology classification using the selected model is also possible.

The following hyperparameters were used in all neural networks:

The Adaptive Moment Estimation (Adam) was chosen as the optimizer (Kingma & Ba, preprint 2014) and the hyperbolic tangent (tanh) as the activation function for the dense hidden layers, while we chose softmax as the activation function for the output layer (Bridle, 1990; Guo et al., 2017). In dropout layers we used a dropout rate of 30%, except for the B10 model where we used 30% and 50% in the different dropout layers. Dropout has been shown to greatly improve the results (Wager et al., 2013). All NNs were trained for 500 epochs, except for the amino acid model selection NNs which were only trained for 50 epochs due to the long training time and a very good convergence after this number of epochs. Categorical-Cross-Entropy was selected as the loss function and accuracy as the validation metric. More details are given in the supplement. Trained NNs can be downloaded with the link given in the Data Availability section below.

### Creating the training data sets

Supervised training of NNs requires extensive training datasets with many thousand alignments for which the correct model and topology are known. Alignments were simulated by starting from a random sequence with nucleotide or amino acid frequencies given by equilibrium frequencies of the substitution model. Sequences evolved under the stochastic process of a given model of sequence evolution along the given tree with branch lengths. In the present work we used the PolyMoSim-v1.1.4 software (https://github.com/cmayer/PolyMoSim) to simulate alignments of four sequences for a given tree and a specified model of sequence evolution for nucleotides or amino acids. PolyMoSim offers a wide range of output formats, including a format that directly outputs pattern frequencies of simulated alignments, which tremendously speeds up the process of creating training data sets.

We generated training data sets for the following prediction tasks:

(I) Prediction of the best fitting nucleotide substitution model for a given four sequence alignment: Models included in the prediction are the JC+I+G (Jukes & Cantor, 1969), K2P+I+G (Kimura, 1980), F81+I+G (Felsenstein, 1981), HKY+I+G (Hasegawa et al., 1985), GTR+I+G (Tavaré, 1986) models. For all models a variable amount of invariant sites (+I) was considered and for all other alignment sites, gamma distributed site rates (+G) were assumed in the training and later in the verification data. The F84 model (Felsenstein & Churchill, 1996) which is included in the topology prediction was not included in the model prediction due to its similarity to the HKY model and its unavailability in the model selection program ModelFinder (Kalyaanamoorthy et al., 2017) implemented in IQ-Tree, version 2.1.3 (Nguyen et al., 2015, Minh et al., 2020), which we used for comparisons.

For each of the five nucleotide substitution models we simulated 90,000 alignments with what we call “normal” internal branch lengths (Table 2) and 90,000 alignments with particularly short internal branch lengths, resulting in a training data set of 900,000 alignments (Table 1). Alignment lengths were 30,000 bp, which is sufficient for sampling the 256 possible site pattern frequencies. The tree topology of each alignment evolved on was chosen randomly among the three possible topologies for four taxa. Branch lengths were drawn randomly from uniform distributions in intervals given in Table 2. Model parameters of the corresponding substitution models were chosen randomly from the distributions and ranges as given in Table 2.

**Table 2:**
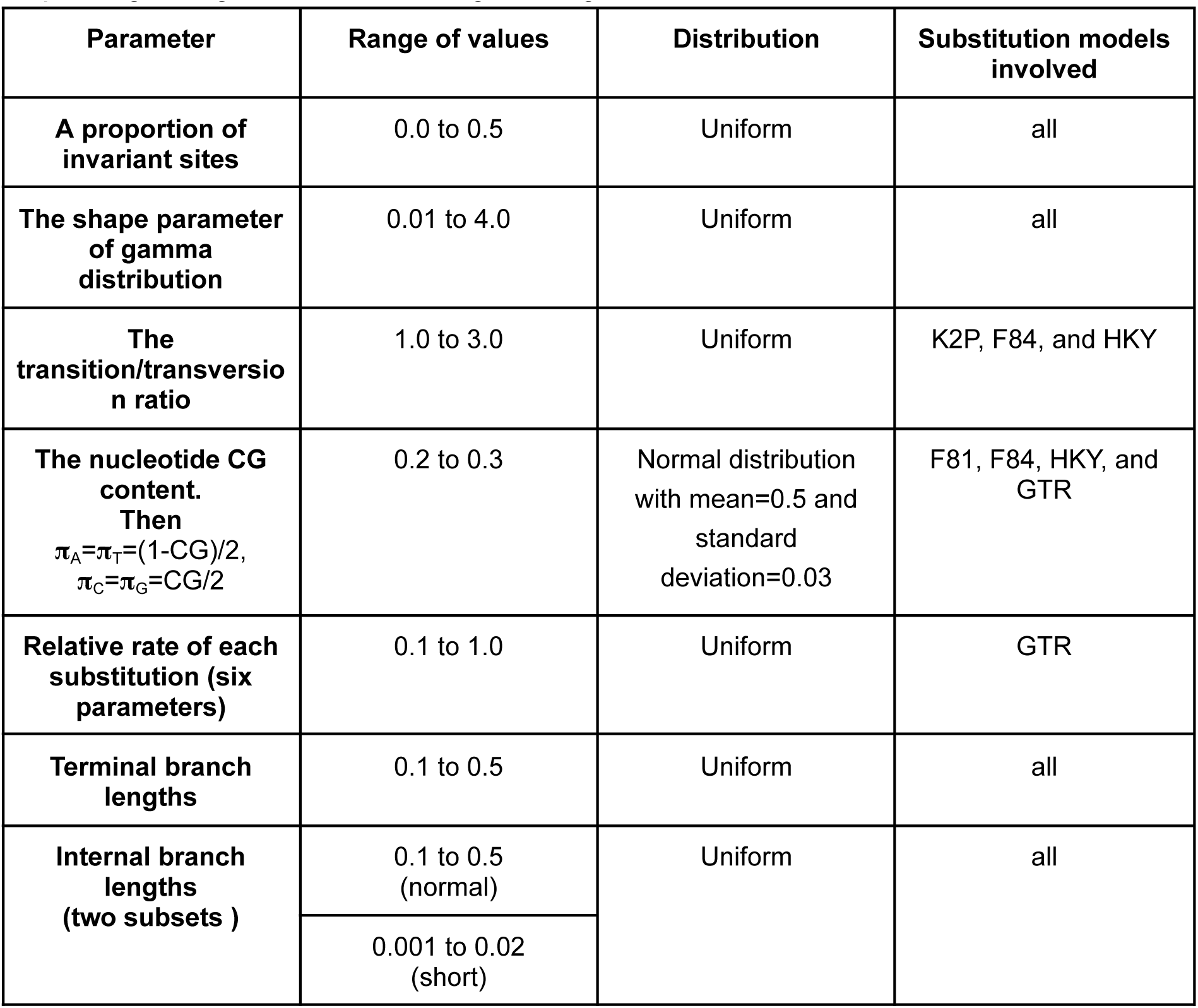
Parameter ranges and distributions used in data simulation. The minimum and maximum values for the proportion of invariant sites, the shape parameter of the gamma distribution and the CG content have been chosen after inspecting a large number of real gene alignments.

(II) Prediction of the best fitting amino acid substitution model for a given four taxon alignment: Models included in the prediction are the JTT+I+G (Jones et al., 1992), LG+I+G (Le & Gascuel, 2008), WAG+I+G (Whelan & Goldman, 2001), Dayhoff+I+G (Dayhoff et al., 1978) models. Again, invariant sites (+I) and a site rate heterogeneity governed by a gamma distribution (+G) have been used in the simulations. Since computing and storing amino acid site pattern frequencies is computationally expensive we decreased the number of simulated data sets to 37,500 per substitution model for normal and the same number for short internal branch lengths, resulting in a training data set of 37,500*4*2=300,000 alignments (Table 1). The sequence length was increased to 1,000,000 aa in order to sample the 160,000 different site patterns more comprehensively. Tree topologies, branch lengths and model parameters were chosen randomly as in the case of the nucleotide model from distributions and ranges given in Table 2.

(III) Prediction of the most likely topology for a given four taxon alignment of nucleotide sequences. As mentioned in the introduction, we trained NNs specific for the nucleotide substitution models (i.e., JC+I+G, K2P+I+G, F81+I+G, F84+I+G, HKY+I+G, GTR+I+G). Each model specific NN was trained with 200,000 alignments of length 100,000 bp for each of the three topologies, i.e.100,000 for normal and short internal branch lengths (see Table 1). Branch lengths and model parameters were chosen randomly from the distributions and ranges given in Table 2.

(IV) Prediction of the most likely topology for a given four taxon alignment of amino acid sequences: Again, we trained NNs specific for the amino acid substitution models (i.e., LG+I+G, JTT+I+G, WAG+I+G, Dayhoff+I+G). Each model specific NN was trained with 20,000 alignments of length 1,000,000 aa for each of the three topologies,i.e., 10,000 for normal and 10,000 short internal branch lengths (see Table 1). Branch lengths and model parameters were chosen randomly from the distributions and ranges given in Table 2.

The number of NNs we trained for each of the prediction tasks are given in Table 1. All trained NNs can be downloaded with the link provided in the Data Availability Section below.

Verification data sets used to test the trained NNs, and to compare prediction accuracies and computation times with other methods/programs have been generated independently but in the same way the training data was generated. Only different alignment lengths have been used in the different comparisons. To verify the substitution model classification, we simulated 1,000 alignments for each substitution model. For the five nucleotide models 5,000 alignments of length 30,000 bp and for the four amino acid models 4,000 alignments of length 10,000 aa were generated. The prediction successes of the NNs were compared to the model prediction of the ModelFinder software implemented in IQ-Tree version 2.1.3. To conduct a fair comparison, the set of models among which ModelFinder chooses the best model was restricted to the models available in the NN prediction by specifying the command line parameter “-mset JC,K2P,F81,HKY,GTR -mrate I,G,I+G” for nucleotide and “-mset JTT,LG,WAG,Dayhoff -mrate I,G,I+G” for amino acid datasets. In ModelFinder the BIC was used to select the best model. A two-tailed binomial test with the standard Wilson score interval and with continuity correction (Wilson, 1927; Crawley, 2014) has been carried out to determine whether the ModelFinder or the NN predictions were significantly better (p-value < 0.05).

To verify the topology prediction, we simulated 1,000 alignments of the lengths 1,000 and 10,000 nucleotide bases and amino acids for each of the three topologies. Two separate verification data sets of the given sizes were created for normal and short internal branch lengths. The topology predictions have been compared with the IQ-Tree implementation of the ML method and the BioNJ method (Gascuel, 1997), an improved implementation of the Neighbour Joining algorithm that is invoked by specifying the -fast option. For the BioNJ tree, corrected distances were determined by conducting a model prediction followed by estimating ML distances. For the ML tree inference, IQ-Tree was not restricted to a set of substitution models but was allowed to select the most appropriate model on its own. Again, the same binomial test was used to determine whether the NN method was significantly better or worse than the ML and/or the BioNJ methods.

### Computing times

Computing times of the NN, ML and BioNj methods have been compared as described in the supplement Section 1.5.

### Alternative machine learning algorithms

Besides NNs, a number of machine learning classifiers exist that are well suited for frequency distribution classifications (see e.g. Hastie et al., 2009). To evaluate the performance of different machine learning algorithms, including the Gaussian process classifier, several implementations of the RF classifier and the SVM classifier, the Lazypredict Python library was utilised (Pandala, 2020). A detailed description of how the alternative machine learning algorithms were trained is given in the supplementary methods section 1.6.

### Data visualisation

Prediction and computation time results for NNs were visualised with the Matplotlib library (Hunter, 2007) using Python3. Lazypredict results were visualised using the Seaborn library (Waskom et al., 2017).

## Results

### Substitution model prediction success

For nucleotide and amino acid alignments, the model prediction accuracy of the best NN architecture B10 is shown in Fig. 3. The mean prediction accuracy of the B10 nucleotide NN was 92.6% which was considerably higher than the prediction accuracy of the ML+BIC based model inference, which showed an accuracy of 87.1%. The NNs were significantly better than the ML method (p-value < 2e-16).

**Fig. 3.**
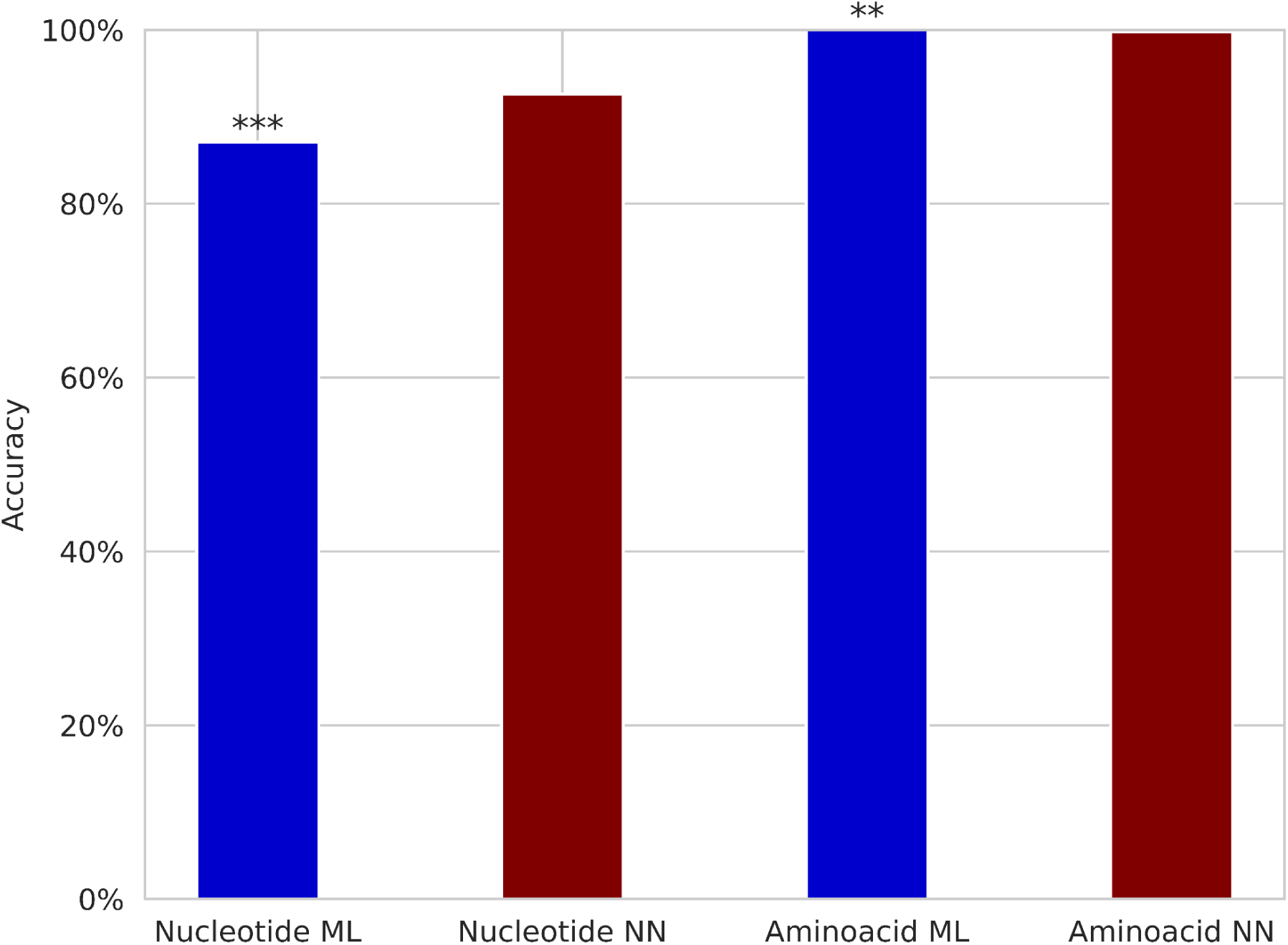
Substitution model prediction accuracy of NNs and ML+BIC on test data sets with alignments of length 30,000 bp and 10,000 aa (with branch lengths ranging from 0.1 to 0.5). p-values with 0.001<p≤0.01 and p≤0.001 are flagged with “**” (medium significance) and “***” (high significance), respectively.

The amino acid NN prediction accuracy of 99.8% obtained with the B10 network is only marginally, but significantly (p-value = 0.0076) worse than the 100% accuracy obtained with ModelFinder for our test conditions. A comparison of the prediction accuracies of different NN architectures for nucleotide and amino acid evolutionary models is presented in supplementary materials Table. S1.

### Topology prediction success

For nucleotide and amino acid alignments, the topology prediction accuracies of the best NN architectures and their comparison with other methods are shown in Figs. 4 to 7. We have trained substitution model specific NNs for the topology prediction, which means that a model has to be selected prior to conducting a topology classification. This is analogous to what we do when using the ML method. Therefore, we tested the NNs with alignments that evolved under the same evolutionary model as the alignments that were used to train the models, but with independently chosen model parameters and branch lengths.

**Fig. 4.**
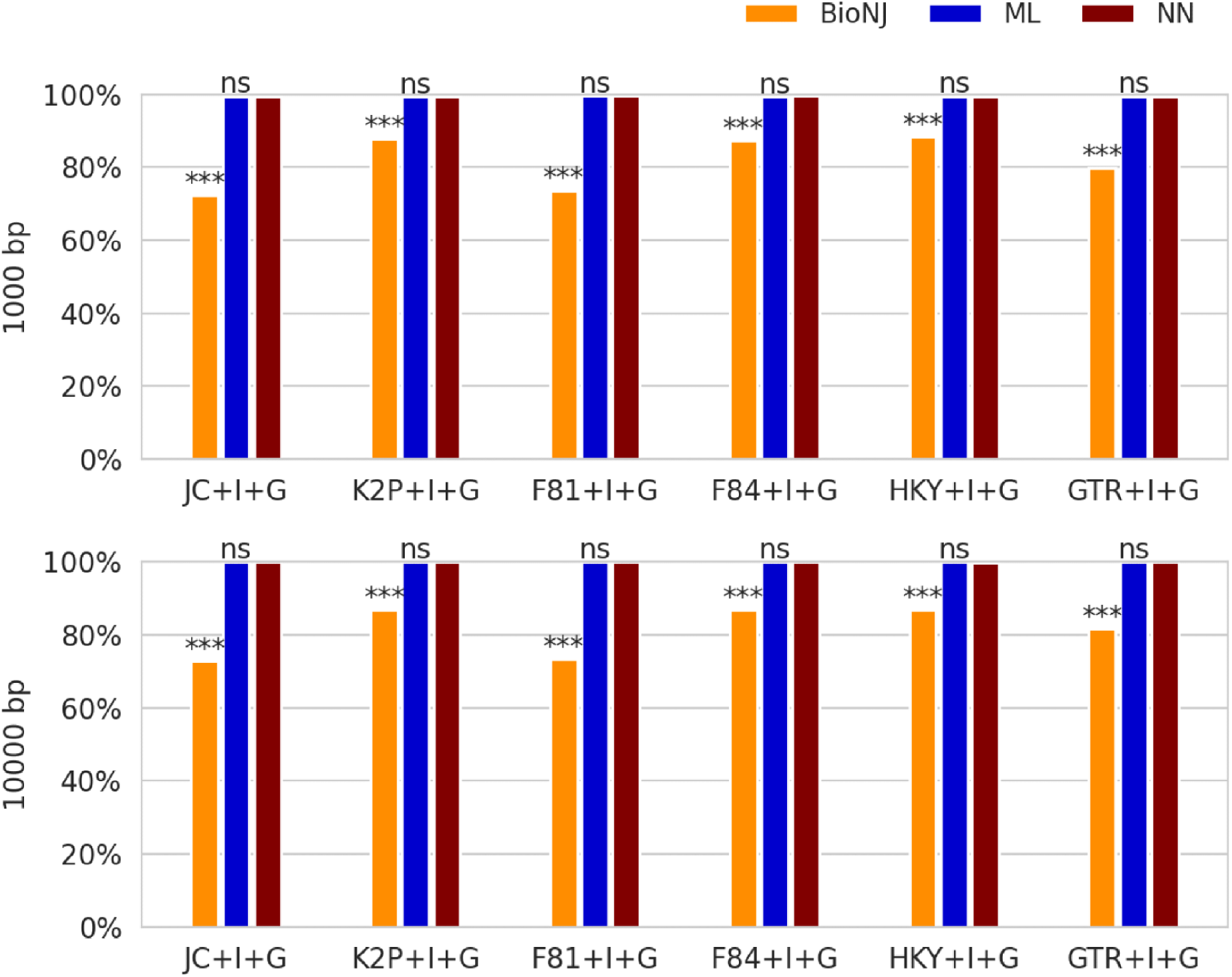
Accuracy of topology reconstruction, with branch lengths ranging from 0.1 to 0.5, using NNs, ML+BIC and BioNJ for 1,000 bp (top figure) and 10,000 bp (bottom figure) long nucleotide alignments. The significance test was conducted between BioNJ and NN and between ML and NN. p-values with p>0.05, and p≤0.001 are flagged with “ns” (non-significant), and “***” (high significance), respectively.

Topology prediction accuracies using NNs, the ML method and the BioNJ method for nucleotide alignments of length 1,000 bp and 10,000 bp and inner branch lengths in the range 0.1<v<0.5 are shown in Fig. 4.

The accuracy of NN topology predictions for nucleotide alignments of length 1,000 bp ranged from 99.4 to 99.6% (Table S.1; Fig. 4). No statistically significant differences were found between NNs and the ML method (p-values ranged from 0.63 to 1.0).

For alignments lengths of 10,000 bp, our NN predictions achieved high accuracies of 99.97% for the HKY model and 100.0% for all other models (see supplementary Table S1 and Fig. 4), which did not differ significantly from the reconstruction success of the ML method (p-value=1.0). In contrast, the BioNJ algorithm showed significantly lower prediction accuracy than the NN classifiers (p-values < 2e-16). A comparison of the six NN architectures showed in supplementary, see Table S1 for details.

The topology reconstruction success of the ML, BioNJ and NN methods were also compared for alignments that were simulated on trees with particularly short internal branches in the range 0.001 < v < 0.02 (Fig. 5). For sequences of length 1,000 bp, we observed a poor accuracy of both, NNs (ranging from 47.6% to 48.3%) and the ML method (ranging form 47.8% and 48.4%), with no significant differences (p-value between 0.74 and 0.94). The BioNJ method’s prediction success rate was 38.4-43.3%, which is better than choosing a topology randomly (33.3%). For alignment lengths of 10,000 bp and for short internal branch lengths the ML method (with accuracies ranging from 71.8% and 73.6%) outperformed the NNs for the K2P+I+G, the F81+I+G model as well as for the F84+I+G model (p-value between 0.01 and 0.04), but not for the other models (p-value between 0.06 and 0.31). Accuracies are given for the best performing NN architecture. A detailed comparison of the accuracies for different NN architectures is given in Supplementary Table S1.

**Fig. 5.**
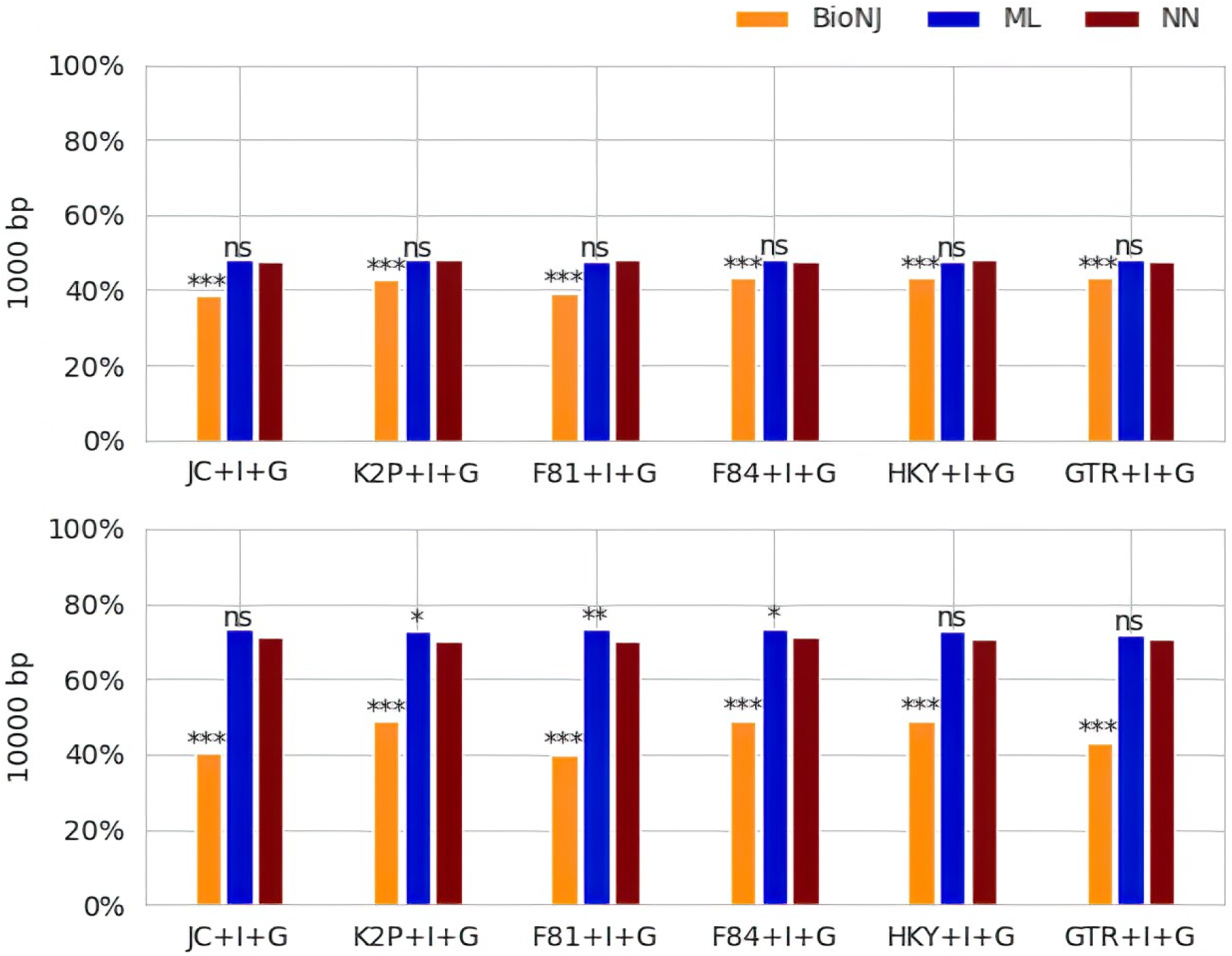
Accuracy of topology reconstruction with internal branch lengths in the range 0.001 to 0.02 using NNs, ML+BIC and BioNJ on 1,000 bp and 10,000 bp long nucleotide alignments. Significance tests were conducted between BioNJ and NN, and between ML and NN. p-values with p > 0.05, 0.01<p≤0.05, 0.001<p≤0.01 and p ≤ 0.001 are flagged with “ns” (non-significant), “*” (low significance), “**” (medium significance), and “***” (high significance), respectively.

For amino acid alignments of length 1,000 aa and 10,000 aa that evolved on trees with an inner branch length in the two ranges 0.1-0.5 and 0.001-0.02, topology prediction success rates are shown in Fig. 6 and Fig. 7, respectively. For normal internal branch lengths and an alignment length of 1,000 aa NNs achieved a prediction accuracy of 97.5-98.3% (Table S1; Fig. 6), while the ML method achieved an accuracy of 99.90-99.97%, showing a significantly better prediction success (p-value <2e-16). The BioNJ method had an accuracy of 91.6-93.0%, which was significantly lower than that of the NNs (p-value < 2e-16). For alignments of length 10,000 amino acid residues, the binomial distribution test did not find any statistically significant difference between NNs and ML method for the JTT+I+G, LG+I+G, and WAG+I+G models (p-value from 0.074 to 0.48), but for the Dayhoff+I+G model (p-value = 0.023). Accuracies achieved with BioNJ were significantly lower for all evolutionary models (p-value < 2e-16).

**Fig. 6.**
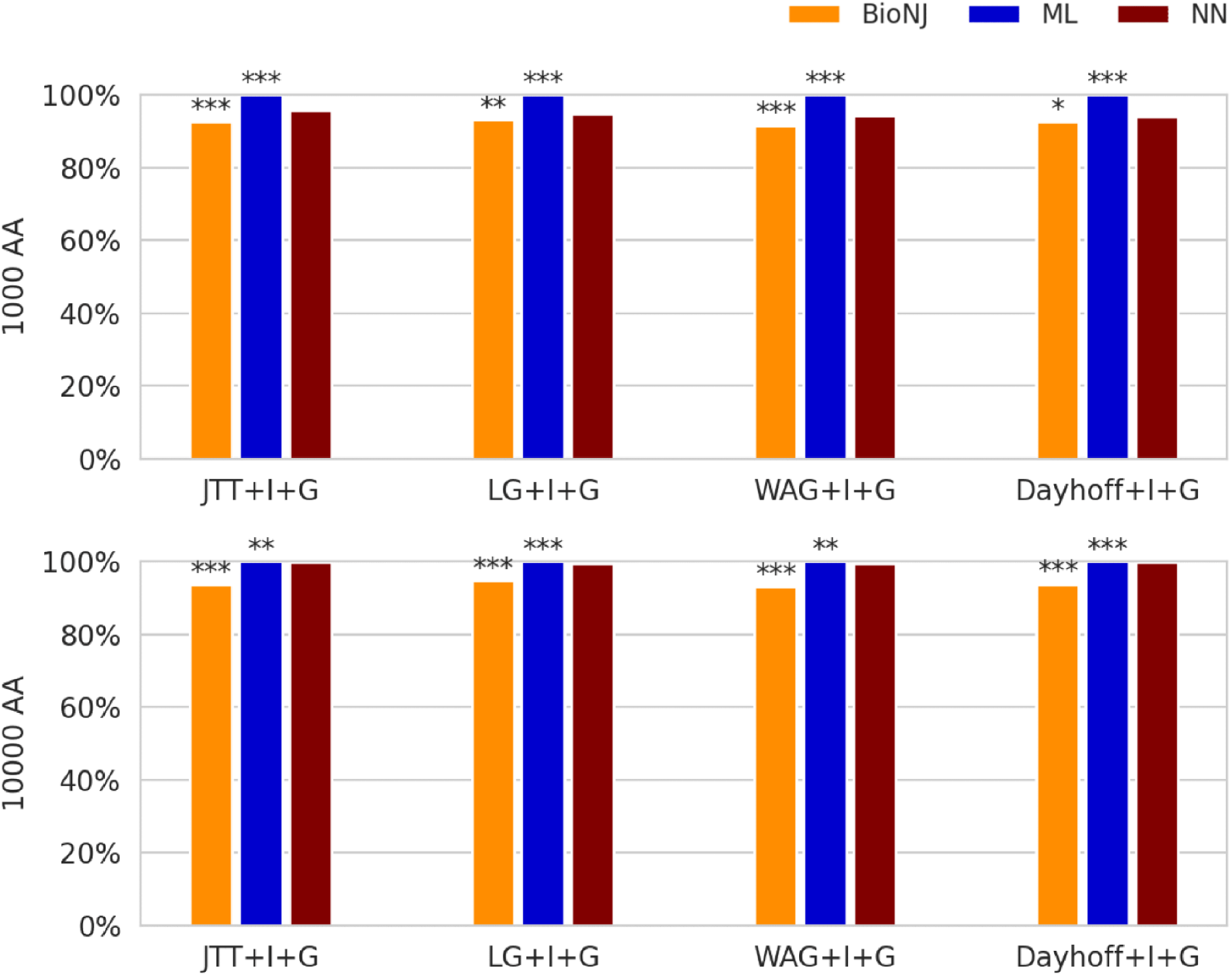
Topology prediction accuracy of topology reconstruction (with internal branch lengths in the range 0.1 to 0.5) using NNs, ML+BIC and BioNJ on 1,000 and 10,000 long amino acid alignments. The results were obtained for the sequences that evolved under four different substitution models: JTT+I+G, LG+I+G, WAG+I+G, Dayhoff+I+G. Significance tests were conducted between BioNJ and NN, and between ML and NN. p-values with p>0.05, 0.01<p≤0.05, 0.001<p≤0.01 and p≤0.001 are flagged with “ns”, “*”, “**”, and “***”, respectively.

**Fig.7.**
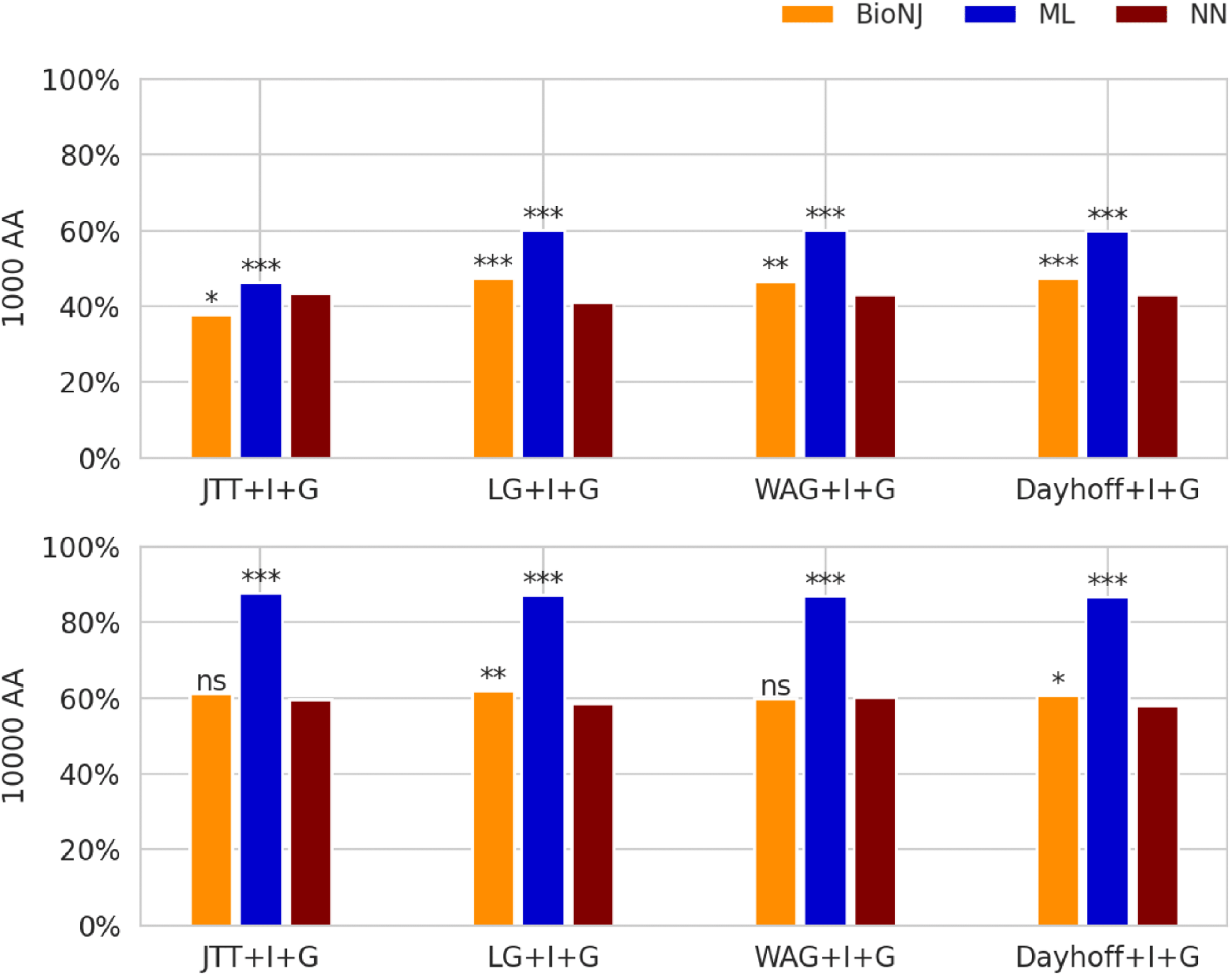
Topology prediction accuracy of topology reconstruction with inner branch lengths ranging from 0.001 to 0.02 using NNs, ML+BIC and BioNJ on 1,000 (top figure) and 10,000 (bottom figure) long amino acid alignments. Significance tests were conducted between NN and BioNJ as well as between NN and ML. p-values with p>0.05, 0.01<p≤0.05, 0.001<p≤0.01 and p ≤ 0.001 are flagged with “ns”, “*”, “**”, and “***”, respectively.

For short internal branch lengths, the NNs achieved an accuracy of 41.2-43.5% (Fig.7). They were significantly outperformed by the ML method (p-value < 2e-16). NNs were only better than the BioNJ method for the JTT model (p-value 0.036). For the other models BioNJ was significantly better than the NNs but with a smaller margin (p-value ranging from 2.6e-06 to 0.0069).

For longer alignments with 10,000 aa, the NNs achieved average accuracies of 58.1-60.2%. They are again outperformed by the ML method (p-value <2e-16) but were as accurate as BioNJ for the JTT and WAG evolutionary models (with p-values of 0.21 and 0.87, respectively), though they remained inferior for the LG (p-value = 0.0056) and Dayhoff (p-value = 0.046) models.

### Alternative machine learning algorithms for topology prediction

For classifiers trained and tested on nucleotide alignments of length 100,000 bp that evolved under the K2P+I+G evolutionary model on trees with normal and short internal branch lengths, we obtained the highest prediction accuracy of 96% with the SVM classifier which is marginally better than the best NN (B2) with 95.4%. Detailed results are provided in the supplementary materials. For amino acid alignments of length 1,000,000 aa that evolved under the Dayhoff+I+G evolutionary model the highest prediction accuracy of 98% was obtained with the LightGBM classifier which is a bit lower than the prediction accuracy of the best NN (B10) with 99.1%. We found that nine out of 25 machine learning algorithms achieved a topology prediction accuracy rate of 90% or higher in predicting the correct tree topology for nucleotide data, whereas 12 out of 26 machine learning algorithms achieved a comparable accuracy for amino acid data sets. See supplementary materials section 2.2 and Figs. S4 for details.

### Prediction times

We found that prediction times for the ML method are at least a factor of 3.5 higher than the prediction times of the NNs for nucleotide alignments of length 1,000 bp and a factor of 16 higher for amino acid alignment of length 1,000 aa. For increasing alignment lengths, the relative advantage of the NNs over the ML method increases. Detailed results are presented in the supplementary results and Figs. S3.

## Discussion and comparison of results

Predicting the model of sequence evolution and the topology which most likely created an alignment is an essential task in molecular biology. Today, the ML method is the gold standard for these inference tasks, since it has been shown that the ML method converges on the correct result as the amount of available data approaches infinity if certain conditions are met. Disadvantages of the ML method are comparatively long computation times as well as a complex procedure to implement new evolutionary scenarios. Alternative methods to the ML method are not only of theoretical interest. They could offer advantages in computational efficiency and facilitate the implementation of more complex evolutionary scenarios in the future.

Our results show that NNs are well suited to predict the best fitting model of sequence evolution for the nucleotide and amino acid models included in our test. For the nucleotide model prediction, NNs were even significantly better than the ModelFinder result using ML+BIC (see supplementary Table S1 for details). A result better than an ML selected model would not be expected and we conjecture that the lower prediction accuracy of ModelFinder is linked to the fact that it does not select the model with the highest likelihood, but instead the model with the best BIC score, which is the most widely used model selection criterion and the default in ModelFinder. A detailed comparison of the NN model predictions with different model selection criteria implemented in ModelFinder would be interesting but is beyond the scope of this paper. For nucleotide alignments, NNs and ModelFinder both yield model prediction accuracies well below 100% which is expected, since many substitution models are nested, i.e. a considerable part of the model parameter space of more complex models can mimic parameter values of a simpler model. For these model parameter combinations, the correct behaviour is to choose the simpler model. In the case of amino acid model predictions, the ML method was marginally but significantly better than the NN (100% vs 99.8%). Since amino acid models are not nested, accuracies close to 100% are achievable. During the course of the project, we have already increased the size of the amino acid training data from a total of 60,000 to 300,000 training alignments, which increased the model prediction accuracy from 88.0% (data not shown) to 99.8%. We expect that an even larger training data set yields a NN that is not significantly worse than ModelFinder. It should be noted that for amino acid alignments of four sequences, we have 160,000 different site patterns. Even for a length of the training sequences of 1,000,000 aa the average number of patterns found per pattern class is only 1,000,000/160,000=6.25. Clearly, this alignment length does not allow us to simulate highly precise pattern frequencies. In order to sample the pattern frequencies sufficiently for all model parameter and branch length combinations, it is expected that (i) longer sequence lengths and (ii) a larger number of training alignments is required. We expect that with more computational resources it should be possible to train amino acid NNs such that they become as accurate as the ML method in predicting the correct substitution model. Indeed, the size of the training data set was limited by the available computational resources required for creating and storing the training data set. The problem is illustrated by the fact that the training data set we generated requires 160,000*300,000*4 Byte=192 GB of computer RAM, if 32-bit floating point numbers are utilised.

Machine learning has also been used by Abadi et al. (2020) and Burgstaller-Muehlbacher et al. (2021) to choose suitable nucleotide substitution models. Abadi et al. (2020) used a RF classifier to choose the model of sequence evolution that yields the best branch length estimates in a phylogenetic tree reconstruction that follows the model selection step. They have shown that their approach is better than the standard model selection criteria to select a substitution model that recovers the correct branch lengths.

Burgstaller-Muehlbacher et al. (2021) analysed images of alignments of 8, 16, 64, 128, 256 and 1024 taxa with CNNs to determine the best model of sequence evolution and to estimate the shape parameter of the gamma distribution. They compared the model prediction accuracies with those of ModelFinder with BIC and found comparable accuracies between the methods. For alignment lengths of 1,000 bp to 100,000 bp they obtained model prediction accuracies that are comparable to the ModelFinder. Since they used branch lengths and model parameters from empirical data sets, the results are difficult to compare with our results since the variance of empirical distributions are smaller than those of the uniform distributions used in this study.

Our results also show that NNs are highly suitable to predict the best tree topology for nucleotide and amino acid alignments. Topology prediction accuracies of NNs for nucleotide alignments were highly similar to those of the ML method. The ML method was marginally but significantly better only for alignments of length 10,000 bp that evolved on a tree with short internal branch lengths (0.001 to 0.02) and for the evolutionary model K2P+I+G, F81+I+G and F84+I+G if the level of significance is used without Bonferroni correction. Since we have conducted multiple tests, some tests are expected to have a p-value <0.05 by chance. With a Bonferroni correction that corrects for multiple tests, the ML method is not significantly better in any of the nucleotide topology predictions. Furthermore, we expect that more training data could remove any difference we have found. It should be noted that short internal branch lengths pose a problem to all tree reconstruction methods, since for an alignment of length of 10,000 bp and a short internal branch length in the range 0.001 to 0.02 we only expect a total of 10-200 substitutions along the inner branch and along the whole alignment. Therefore, the site pattern frequencies of alignments that evolved on different topologies will differ only marginally if the inner branch length is very short. The high number of misclassifications of the ML and NN method for short internal branch lengths are most likely attributed to cases in which site pattern frequencies of alignments favour the wrong topology by chance. In this case only longer sequences can help to reconstruct the correct topology.

Suvorov (2020) showed that CNNs which analyse images of alignments can be as good as ML and other methods for alignments that evolved on trees in the so-called Farris- and Felsenstein zones. For alignments of length 1,000 bp that evolved on trees with internal branches in the range 0 to 0.5, their CNNs showed an accuracy of 82%, which is as accurate as the ML method (82%) for the same dataset. In our study, NNs showed accuracies above 99% for internal branches in the range 0.1 to 0.5 for the same alignment length. For alignments of length 1,000 bp that evolved on trees with an internal branch length in the range of 0.0 to 0.05, their CNNs achieved an accuracy of 52%, while the ML method achieved 57%. Our NNs also showed accuracies around 48%, with internal branch lengths in the range 0.001 to 0.02. Due to the different parameter and branch length ranges used when simulating the alignments, only an approximate comparison of the accuracies is possible. Also Zou et al. (2020) proposed CNNs to analyse images of alignments CNNs that showed accuracies on quartet trees that are comparable to the ML and Bayesian methods. Their approach showed accuracies in the range from 90% to 98% when alignment lengths increased from 100 to 10,000 bp. The branch lengths were in the range from 0.02 to 2 which implies a larger mean branch length than in our simulations and leads to higher accuracies in their reconstructions.

For amino acid alignments the ML method was always significantly better than NNs for alignments of the length 1,000 aa and for alignments generated with short internal branch lengths. For alignment lengths of 10,000 aa and normal internal branch lengths, the differences between ML and NNs would not be significant if the level of significance would be adjusted for multiple testing using the Bonferroni correction procedure. As in the case of the amino acid model selection, the size of the training data set was limited by the available computational resources. Our initial training data set was even smaller with only 6,000 training alignments of length 1,000,000 aa. For the final amino acid NNs we used 60,000 training alignments of length 1,000,000 aa. When testing the trained NNs with alignments of length 1,000 aa and normal internal branch lengths the prediction accuracy for the four amino acid models improved considerably from the range 94.0 to 95.5% (data not shown) to a range of 97.5 to 98.3% (Table S1). This suggests that by further increasing the size of the training data set, it is possible to achieve higher accuracies. Altogether we have found prediction accuracies as good as or almost as good as the prediction accuracies obtained with the ML method.

Site pattern frequencies have previously been used by Leuchtenberger et al. (2020) to choose the best tree reconstruction method and by Suvorov & Schrider (preprint 2022) to predict branch lengths. In contrast, Zou et al. (2020) and Suvorov et al. (2020) suggested analysing images of alignments with CNNs to predict the best tree topology. Due to the width of the convolutional kernel, this approach combines the information about neighbouring alignment sites, which are consequently not treated as independent and identically distributed. It is possible that this extracts information about the co-evolution of neighbouring alignment sites from an alignment, but the influence of this approach has not been investigated yet. Presumably, these NNs would be specific for genomic regions since their convolutional kernels will include information about co-evolving neighbouring alignment sites with the feature or drawback that different NNs would need to be trained for different genomic regions. A classification with site pattern frequencies will not include information about co-evolving neighbouring alignment sites.

A problem we faced during training of NNs was the need to normalise the site pattern frequencies before passing them to the first dense layer for some of the prediction tasks. Even though the site pattern frequencies are all numbers in the range of 0 to 1, we found that a normalisation is indispensable for some of the NNs. In particular the amino acid NNs often did not train at all if non-normalised data was used, resulting in training accuracies below 50% in the limit of long training times or in an oscillation of the training accuracies. With normalisation training and prediction data, this problem did not occur. The methods we used to normalise the data is described in detail in supplementary materials section 1.2.

In contrast to the ML method which evaluates an optimality criterion, the machine learning methods learn the site pattern frequency distributions that are associated with certain substitution models and tree topologies. By training a classifier, one circumvents optimising the likelihood function, which can be computationally more efficient. In order for a machine learning method to be able to predict the correct model and topology, the training data must sample site pattern frequencies sufficiently dense for all combinations of the substitution model parameters and branch lengths that shall be classified, in particular for the boundary cases. If NNs are used for the classification, they have to be designed such that they are sufficiently complex to be able to “memorise” the frequency distributions they were trained with. For this, the network architecture, the number of layers and neurons as well as other hyperparameters, such as the activation function and the optimizer have to be optimised for the specific classification task. In this project we tried to optimise the number of layers, the number of neurons and other hyperparameters such as the activation function, the optimiser and the dropout rate for all six architectures by conducting grid searches for selected parameters. Since the number of possible hyperparameter combinations is huge, it is possible that other hyperparameter combinations lead to even higher prediction accuracies.

In this study we compared six NN architectures ranging from a simple sequential NN to NNs with multiple branches and a NN with interconnections. The NNs with more neurons and more trainable parameters are also able to memorise more information. For some of the classification tasks the best NN architecture is significantly better than all other architectures (see supplementary Table S1 for details). Altogether, the B10 architecture was the best for both substitution model prediction tasks. For most topology predictions, the B3 architecture is preferred and for the few cases for which B3 is not the best architecture, it was not significantly worse. Therefore, the NN architecture implemented in B3 is a good starting point when designing NNs for topology predictions. The most complex architecture CU was the best architecture for several topology prediction tasks, but it could not outperform the other architectures significantly. While we have a significant difference in the number of features used in the nucleotide and amino acid data sets, we see no general trend that the simpler architectures are better suited for nucleotide data or more complex architectures are better suited for amino acid data.

We have also compared the NN classifiers with alternative machine learning classifiers such as SVM and RF classifiers. For selected test cases we found that the best performing alternative classifiers are as good or almost as good as the best NNs. Certainly, these classifiers should be explored in more detail in future studies. Their main advantages are faster training and predictions and that the trained models have much smaller file sizes and would be much easier to use.

Currently, the main disadvantage of using machine learning methods for phylogenetic tree reconstruction is that these methods are difficult to extend beyond a few taxa. The reason is that classifiers have to be trained with a large number of data sets for each possible topology, which quickly becomes infeasible as the number of taxa increases. Nevertheless, we think that our NN classifiers will be useful in future research.

The main advantage of utilising the machine learning approaches proposed here is their increased speed in determining the best model of sequence evolution and topology compared to ML method. For short nucleotide alignments, the NNs are about 3.5 times faster than the ML method and the advantage of the NNs increases for longer alignments and amino acid sequences. We have mentioned in the supplement that an even faster prediction would be possible if multiple predictions are combined or if a GPU is used.

Despite the fact that NN classifiers are currently limited to four taxa, there are a number of potential applications. Beside their computational advantage, the following applications are conceivable:

(i) The PartitionFinder program (Lanfear et al., 2012; Lanfear et al., 2017) spends most of its computing time on determining the best model of sequence evolution for a large number of subsets of a data set. Using NNs could tremendously speed up the process of finding meta-partitions that best evolve under the same substitution model. (ii) Massive computations of quartet trees are used e.g. in approaches such as Quartet Puzzling (Strimmer & von Haeseler, 1996), Likelihood mapping (Strimmer & von Haeseler, 1997), and Sliding window phylogenetic analyses to detect introgression (see e.g. Hibbins & Hahn, 2022). These methods could benefit from a fast topology prediction method for four sequences.

## Conclusion

In this study we have shown that NNs trained on site pattern frequencies of four taxon alignments are as accurate or close to being as accurate as the ML method to select the best fitting nucleotide or amino acid evolutionary model as well as the tree topology on which the sequences in the alignment evolved. In cases NNs are not as accurate as the ML method, we provide evidence that a larger training data set should be able to close the accuracy gap. The main advantage of NNs is that they are much faster than the ML method. The speed difference ranges from a factor of about 3.5 on short nucleotide alignments to more than 1000 for the long amino acid alignments analysed in this study. Furthermore, we have shown that other machine learning classifiers such as SVM and the LGBM classifier are about as accurate as the NNs.

## Data availability

The software for the NNs implementation as well as scripts that have been used in this study can be found at GitHub (https://github.com/cmayer/DeepNNPhylogeny). The PolyMoSim-v1.1.4 program that is needed for training data generation can be obtained at (https://github.com/cmayer/PolyMoSim). The trained machine learning models can be downloaded from the DryAd repository with the DOI https://doi.org/10.5061/dryad.ksn02v783.

## Author contributions

CM conceptualised the project with major contributions from NK. FD and CM designed the initial neural networks. NK conducted most of the research including the final implementation and evaluation of the machine learning concepts and the statistical analyses. NK and CM drafted the manuscript. All authors contributed to and approved the final manuscript.

## Acknowledgement

We cordially thank Marie Brasseur for valuable suggestions on the manuscript. Furthermore, we are thankful to the anonymous reviewers for their helpful comments.

