## Supplementary Material for "Machine learning can be as good as maximum likelihood when reconstructing phylogenetic trees and determining the best evolutionary model on four taxon alignments" (from another version of this preprint: version 1, 2023-07-13)

### 1. Supplementary Methods

#### 1.1 Description of neural network

Our machine learning methods, including the training and prediction scripts, are implemented in the Python programming language. They make use of the Tensorflow (Abadi et al., 2016) and Keras (Chollet et al., 2015) libraries for implementing neural networks, the Scikit-learn library (Pedregosa et al., 2011) for data preprocessing and testing alternative machine learning classifiers and the Numpy library (Harris et al., 2020) for handling large data arrays.

We designed six neural network architectures to conduct substitution model and topology predictions for given four sequence nucleotide and amino acid alignments. These architectures range from a simple sequential network of dense layers to a U-shaped neural network with inner connections. The number of neurons in the output layer varied depending on the classification task. We conducted a considerable search to optimise the neural networks and other hyperparameters. Further improvements could potentially be achieved due to the substantial range of hyperparameter combinations that are possible.

The code for these networks as well as the code for the training and prediction programs is provided in the DeepNNPhylogeny github repository. Trained neural networks can be found in the DryAd repository https://doi.org/10.5061/dryad.ksn02v783. Custom neural networks can be trained by researchers using code from the github page of DeepNNPhylogeny. We refer to the six neural network architectures as B1, B2, B3, B10, U, and CU. The code for these architectures is given in Supplementary materials Section 1.3 below. Graph representations of the neural networks are shown in Figures S1 and S2.

With Input-LEN we want to denote the number of neurons in the input layer. This number corresponds to the number of different site patterns that are possible in four sequence alignments. For nucleotide data Input-LEN=$4^4$=256 and for amino acid data Input-LEN=$20^4$=160,000. Output-LEN is the number of neurons in the output layer. This number is equal to the number of classes of the corresponding classification problem. We have Output-LEN = 3 in the case of topology predictions, Output-LEN=5 for nucleotide substitution model predictions (JC, K2P, F81, HKY, GTR) and Output-LEN=4 for amino acid substitution model predictions (Dayhoff, JTT, WAG, LG).

#### 1.2 Input data normalisation

We have tested different forms of data normalisation after we found out that not all neural networks train well if the input data is not normalised. Especially some amino acid neural networks converged to training accuracies below 50% or started to oscillate if the input data was not normalised. Two normalisation methods worked equally well:
Using (a) the `sklearn.preprocessing.StandardScaler` class and (b) the `tf.keras.layers.Normalization` layer class. Since the fitted StandardScaler object of

the former method cannot easily be saved and deployed with the neural networks, we decided to include a normalisation layer for which the trained normalisation parameters are saved together with the weights and biases of the neural network.
For the DNA substitution model prediction the neural network with normalisation layer was slightly but significantly less accurate compared to the model without any normalisation. For this single case we did not use a normalisation layer and provided the trained models without normalisation.

#### 1.3 Source code of neural networks used in this study

As described in the main text, the neural networks obtain site pattern frequencies and the corresponding correct classification as training data. Neural networks are created by the model creating functions defined below. Before calling these functions, we create a single normalisation layer object and train this layer with the full training data stored in the `frequency_train` array.

```
normlayer = tf.keras.layers.experimental.preprocessing.Normalization()
normlayer.adapt(frequency_train)
```

This layer is passed as a parameter to all functions creating the neural networks and it is included as the first hidden layer in all neural networks. Its task is to normalise the training and prediction data before it is passed to the classification layers. The layer is trained once and the parameter values needed for the normalisation are stored together with the neural network.

##### The one branch model (B1)

The B1 model is a simple sequential model with a series of dense and dropout layers. See also Figure S1.

```
def branched_model_1(normlayer):
  input = tf.keras.Input(shape=(Input-LEN,))
  inputn = normlayer(input)

  x = tf.keras.layers.Dense(units=256, activation="tanh")(inputn)
  x = tf.keras.layers.Dropout(0.3)(x)
  x = tf.keras.layers.Dense(units=256, activation="tanh")(x)
  x = tf.keras.layers.Dropout(0.3)(x)
  x = tf.keras.layers.Dense(units=256, activation="tanh")(x)
  x = tf.keras.layers.Dense(units=128, activation="tanh")(x)

  y = tf.keras.layers.Dense(units=64, activation="tanh")(x)
  y = tf.keras.layers.Dropout(0.3)(y)
  y = tf.keras.layers.Dense(units=32, activation="tanh")(y)
  y = tf.keras.layers.Dense(units=Output-LEN, activation='softmax')(y)
  ann = tf.keras.Model(inputs=[input], outputs=y)
  ann.compile(optimizer=”adam”, loss='categorical_crossentropy', metrics=['accuracy'])
  return ann
```

**Two branch model (B2)**

In the B2 model we have two separate paths from the input-layer to the output-layer. Each path consists of a series of dense and dropout layers. See also Figure S1.

```
def branched_model_2(normlayer):
    input = tf.keras.Input(shape=(Input-LEN,))
    inputn = normlayer(input)

    x = tf.keras.layers.Dense(units=256, activation="tanh")(inputn)
    x = tf.keras.layers.Dropout(0.3)(x)
    x = tf.keras.layers.Dense(units=256, activation="tanh")(x)
    x = tf.keras.layers.Dropout(0.3)(x)
    x = tf.keras.layers.Dense(units=256, activation="tanh")(x)
    x = tf.keras.layers.Dense(units=128, activation="tanh")(x)

    x_ann = tf.keras.Model(inputs=input, outputs=x)

    y = tf.keras.layers.Dense(units=256, activation="tanh")(inputn)
    y = tf.keras.layers.Dropout(0.3)(y)
    y = tf.keras.layers.Dense(units=128, activation="tanh")(y)
    y = tf.keras.layers.Dropout(0.3)(y)
    y = tf.keras.layers.Dense(units=128, activation="tanh")(y)

    y_ann = tf.keras.Model(inputs=input, outputs=y)

    combined = tf.keras.layers.concatenate([x_ann.output, y_ann.output])

    z = tf.keras.layers.Dense(units=128, activation="tanh")(combined)
    z = tf.keras.layers.Dropout(0.3)(z)
    z = tf.keras.layers.Dense(units=64, activation="tanh")(combined)
    z = tf.keras.layers.Dropout(0.3)(z)
    z = tf.keras.layers.Dense(units=32, activation="tanh")(z)
    z = tf.keras.layers.Dense(units=Output-LEN, activation='softmax')(z)
    ann = tf.keras.Model(inputs=[input], outputs=z)
    ann.compile(optimizer=”adam”, loss='categorical_crossentropy', metrics=['accuracy'])
    return ann
```

**Three branch model (B3)**

In the B3 model we have three separate paths from the input-layer to the output-layer. Each path consists of a series of dense and dropout layers.

```
def branched_model_3(normlayer):
    input = tf.keras.Input(shape=(Input-LEN,))
    inputn = normlayer(input)

    x = tf.keras.layers.Dense(units=256, activation="tanh")(inputn)
    x = tf.keras.layers.Dropout(0.3)(x)
    x = tf.keras.layers.Dense(units=256, activation="tanh")(x)
```

```
    x = tf.keras.layers.Dropout(0.3)(x)
    x = tf.keras.layers.Dense(units=128, activation="tanh")(x)

    x_ann = tf.keras.Model(inputs=input, outputs=x)

    y = tf.keras.layers.Dense(units=256, activation="tanh")(inputn)
    y = tf.keras.layers.Dropout(0.3)(y)
    y = tf.keras.layers.Dense(units=256, activation="tanh")(y)
    y = tf.keras.layers.Dropout(0.3)(y)
    y = tf.keras.layers.Dense(units=128, activation="tanh")(y)

    y_ann = tf.keras.Model(inputs=input, outputs=y)

    w = tf.keras.layers.Dense(units=256, activation="tanh")(inputn)
    w = tf.keras.layers.Dropout(0.3)(w)
    w = tf.keras.layers.Dense(units=128, activation="tanh")(w)
    w = tf.keras.layers.Dropout(0.3)(w)
    w = tf.keras.layers.Dense(units=128, activation="tanh")(w)
    w_ann = tf.keras.Model(inputs=input, outputs=w)

    combined = tf.keras.layers.concatenate([x_ann.output, y_ann.output, w_ann.output])

    z = tf.keras.layers.Dense(units=256, activation="tanh")(combined)
    z = tf.keras.layers.Dropout(0.3)(z)
    z = tf.keras.layers.Dense(units=128, activation="tanh")(z)
    z = tf.keras.layers.Dropout(0.3)(z)
    z = tf.keras.layers.Dense(units=64, activation="tanh")(z)
    z = tf.keras.layers.Dense(units=Output-LEN, activation='softmax')(z)
    ann = tf.keras.Model(inputs=[input], outputs=z)
    ann.compile(optimizer="adam", loss='categorical_crossentropy', metrics=['accuracy'])
    return ann
```

**Ten branch model (B10)**

In the B10 model we have ten separate paths from the input-layer to the output-layer. Each path consists of a series of dense and dropout layers.

```
def branched_model_10(normlayer):
    input = tf.keras.Input(shape=(Input-LEN,))
    inputn = normlayer(input)

    x1 = tf.keras.layers.Dense(units=256, activation="tanh")(inputn)
    x1 = tf.keras.layers.Dropout(0.3)(x1)
    x1 = tf.keras.layers.Dense(units=128, activation="tanh")(x1)
    x1 = tf.keras.layers.Dropout(0.3)(x1)
    x1 = tf.keras.layers.Dense(units=64, activation="tanh")(x1)
    x1 = tf.keras.layers.Dropout(0.3)(x1)
    x1 = tf.keras.layers.Dense(units=32, activation="tanh")(x1)
    x1_ann = tf.keras.Model(inputs=input, outputs=x1)

    x2 = tf.keras.layers.Dense(units=256, activation="tanh")(inputn)
```

```
x2 = tf.keras.layers.Dropout(0.3)(x2)
x2 = tf.keras.layers.Dense(units=128, activation="tanh")(x2)
x2 = tf.keras.layers.Dropout(0.3)(x2)
x2 = tf.keras.layers.Dense(units=32, activation="tanh")(x2)
x2_ann = tf.keras.Model(inputs=input, outputs=x2)

x3 = tf.keras.layers.Dense(units=256, activation="tanh")(inputn)
x3 = tf.keras.layers.Dropout(0.3)(x3)
x3 = tf.keras.layers.Dense(units=32, activation="tanh")(x3)
x3_ann = tf.keras.Model(inputs=input, outputs=x3)

x4 = tf.keras.layers.Dense(units=256, activation="tanh")(inputn)
x4 = tf.keras.layers.Dropout(0.5)(x4)
x4 = tf.keras.layers.Dense(units=16, activation="tanh")(x4)
x4 = tf.keras.layers.Dropout(0.5)(x4)
x4 = tf.keras.layers.Dense(units=32, activation="tanh")(x4)
x4_ann = tf.keras.Model(inputs=input, outputs=x4)

x5 = tf.keras.layers.Dense(units=32, activation="tanh")(inputn)
x5 = tf.keras.layers.Dropout(0.3)(x5)
x5 = tf.keras.layers.Dense(units=32, activation="tanh")(x5)
x5_ann = tf.keras.Model(inputs=input, outputs=x5)

x6 = tf.keras.layers.Dense(units=64, activation="tanh")(inputn)
x6 = tf.keras.layers.Dropout(0.3)(x6)
x6 = tf.keras.layers.Dense(units=32, activation="tanh")(x6)
x6_ann = tf.keras.Model(inputs=input, outputs=x6)

x7 = tf.keras.layers.Dense(units=128, activation="tanh")(inputn)
x7 = tf.keras.layers.Dropout(0.5)(x7)
x7 = tf.keras.layers.Dense(units=32, activation="tanh")(x7)
x7_ann = tf.keras.Model(inputs=input, outputs=x7)

x8 = tf.keras.layers.Dense(units=111, activation="tanh")(inputn)
x8 = tf.keras.layers.Dropout(0.3)(x8)
x8 = tf.keras.layers.Dense(units=32, activation="tanh")(x8)
x8_ann = tf.keras.Model(inputs=input, outputs=x8)

x9 = tf.keras.layers.Dense(units=77, activation="relu")(inputn)
x9 = tf.keras.layers.Dropout(0.3)(x9)
x9 = tf.keras.layers.Dense(units=32, activation="relu")(x9)
x9_ann = tf.keras.Model(inputs=input, outputs=x9)

x10 = tf.keras.layers.Dense(units=155, activation="relu")(inputn)
x10 = tf.keras.layers.Dropout(0.3)(x10)
x10 = tf.keras.layers.Dense(units=64, activation="relu")(x10)
x10 = tf.keras.layers.Dropout(0.3)(x10)
x10 = tf.keras.layers.Dense(units=32, activation="relu")(x10)
x10_ann = tf.keras.Model(inputs=input, outputs=x10)

combined = tf.keras.layers.concatenate(
    [x1_ann.output, x2_ann.output, x3_ann.output, x4_ann.output, x5_ann.output,
```

```
    x6_ann.output, x7_ann.output, x8_ann.output, x9_ann.output, x10_ann.output])

  z = tf.keras.layers.Dense(units=256, activation="tanh")(combined)
  z = tf.keras.layers.Dropout(0.3)(z)
  z = tf.keras.layers.Dense(units=64, activation="tanh")(z)
  z = tf.keras.layers.Dropout(0.3)(z)
  z = tf.keras.layers.Dense(units=32, activation="tanh")(z)
  z = tf.keras.layers.Dense(units=Output-LEN, activation='softmax')(z)
  ann = tf.keras.Model(inputs=[input], outputs=z)
  ann.compile(optimizer=”adam”, loss='categorical_crossentropy', metrics=['accuracy'])
  return ann
```

**U-model (U)**

The U model is a sequential model with first a decreasing number of neurons per layer (descending layers) followed by an increasing number of neurons per layer (ascending layers) and a final classification step with a decreasing number of neurons (Figure S2). It has the same number of neurons in the different layers as the connected CU-model below and was introduced to allow a comparison between the models with and without interconnections.

```
def branched_model_1u(normlayer):
  input = tf.keras.Input(shape=(Input-LEN,))
  inputn = normlayer(input)

  x3 = tf.keras.layers.Dense(units=256, activation="tanh")(
    inputn)
  x4 = tf.keras.layers.Dropout(0.3)(x3)
  x5 = tf.keras.layers.Dense(units=128, activation="tanh")(x4)
  x6d = tf.keras.layers.Dropout(0.3)(x5)
  x7 = tf.keras.layers.Dense(units=64, activation="tanh")(x6d)
  x7d = tf.keras.layers.Dropout(0.3)(x7)
  x8 = tf.keras.layers.Dense(units=64, activation="tanh")(x7d)
  x9 = tf.keras.layers.Dense(units=128, activation="tanh")(x8)

  x10 = tf.keras.layers.Dense(units=256, activation="tanh")(x9)

  x11 = tf.keras.layers.Dense(units=256, activation="tanh")(x10)
  x14 = tf.keras.layers.Dropout(0.3)(x11)
  x15 = tf.keras.layers.Dense(units=128, activation="tanh")(x14)
  x16 = tf.keras.layers.Dropout(0.3)(x15)
  x17 = tf.keras.layers.Dense(units=64, activation="tanh")(x16)
  x18 = tf.keras.layers.Dense(units=32, activation="tanh")(x17)
  x19 = tf.keras.layers.Dense(units=16, activation="tanh")(x18)
  x20 = tf.keras.layers.Dense(units=Output-LEN, activation="softmax")(x19)

  ann = tf.keras.Model(inputs=[input], outputs=x20)
  ann.compile(optimizer=”adam”, loss='categorical_crossentropy', metrics=['accuracy'])

  return ann
```

**CU-model (CU)**

In the CU-model (connected U-shaped model) we have a decreasing number of neurons in the left branch of the U and an increasing number of neurons in the right branch of the U. There are three horizontal connections within the CU-model (Figure S2). While the CU-model architecture is distantly similar to the U-Net proposed for image segmentation by Ronneberger et al. (2015), we simply wanted to introduce a non-trivial architecture with a large number of neurons that can be trained fast in a stable manner. The ascending branch has no direct application as in the U-Net. The main differences of the CU and the classical U-Net are: (i) The CU architecture is not a transformer model, but used for classification, (ii) it consists of dense and not convolutional layers. The common features are the descending branch followed by an ascending branch, as well as the interconnection between the branches. The U and CU models have the same number and order of layers, and the same number of nodes in the layer. The difference between the U and CU model is that the latter has three interconnections.

```
def branched_model_cu(normlayer):
    input = tf.keras.Input(shape=(Input-LEN,))
    inputn = normlayer(input)

    x3 = tf.keras.layers.Dense(units=256, activation="tanh")(inputn)
    x4 = tf.keras.layers.Dropout(0.3)(x3)
    x5 = tf.keras.layers.Dense(units=128, activation="tanh")(x4)
    x6d = tf.keras.layers.Dropout(0.3)(x5)
    x7 = tf.keras.layers.Dense(units=64, activation="tanh")(x6d)
    x7d = tf.keras.layers.Dropout(0.3)(x7)
    x8 = tf.keras.layers.Dense(units=64, activation="tanh")(x7d)
    y0 = tf.keras.layers.concatenate([x7, x8])

    x9 = tf.keras.layers.Dense(units=128, activation="tanh")(y0)
    y1 = tf.keras.layers.concatenate([x5, x9])

    x10 = tf.keras.layers.Dense(units=256, activation="tanh")(y1)
    y2 = tf.keras.layers.concatenate([x10, x3])

    x11 = tf.keras.layers.Dense(units=256, activation="tanh")(y2)
    x14 = tf.keras.layers.Dropout(0.3)(x11)
    x15 = tf.keras.layers.Dense(units=128, activation="tanh")(x14)
    x16 = tf.keras.layers.Dropout(0.3)(x15)
    x17 = tf.keras.layers.Dense(units=64, activation="tanh")(x16)
    x19 = tf.keras.layers.Dense(units=16, activation="tanh")(x17)
    x20 = tf.keras.layers.Dense(units=Output-LEN, activation="softmax")(x19)

    ann = tf.keras.Model(inputs=[input], outputs=x20)
    ann.compile(optimizer="adam", loss='categorical_crossentropy', metrics=['accuracy'])
    return ann
```

#### 1.4 Graphs of neural networks used in this study

Figures S1 and S2 show the graphical representation of the neural networks B1, B2, U and CU. The networks B3 and B10 are not shown since their graphs do not fit on one page without making the font too small. These images are available as PNG files in the github repository of [DeepNNPhylogeny](DeepNNPhylogeny).

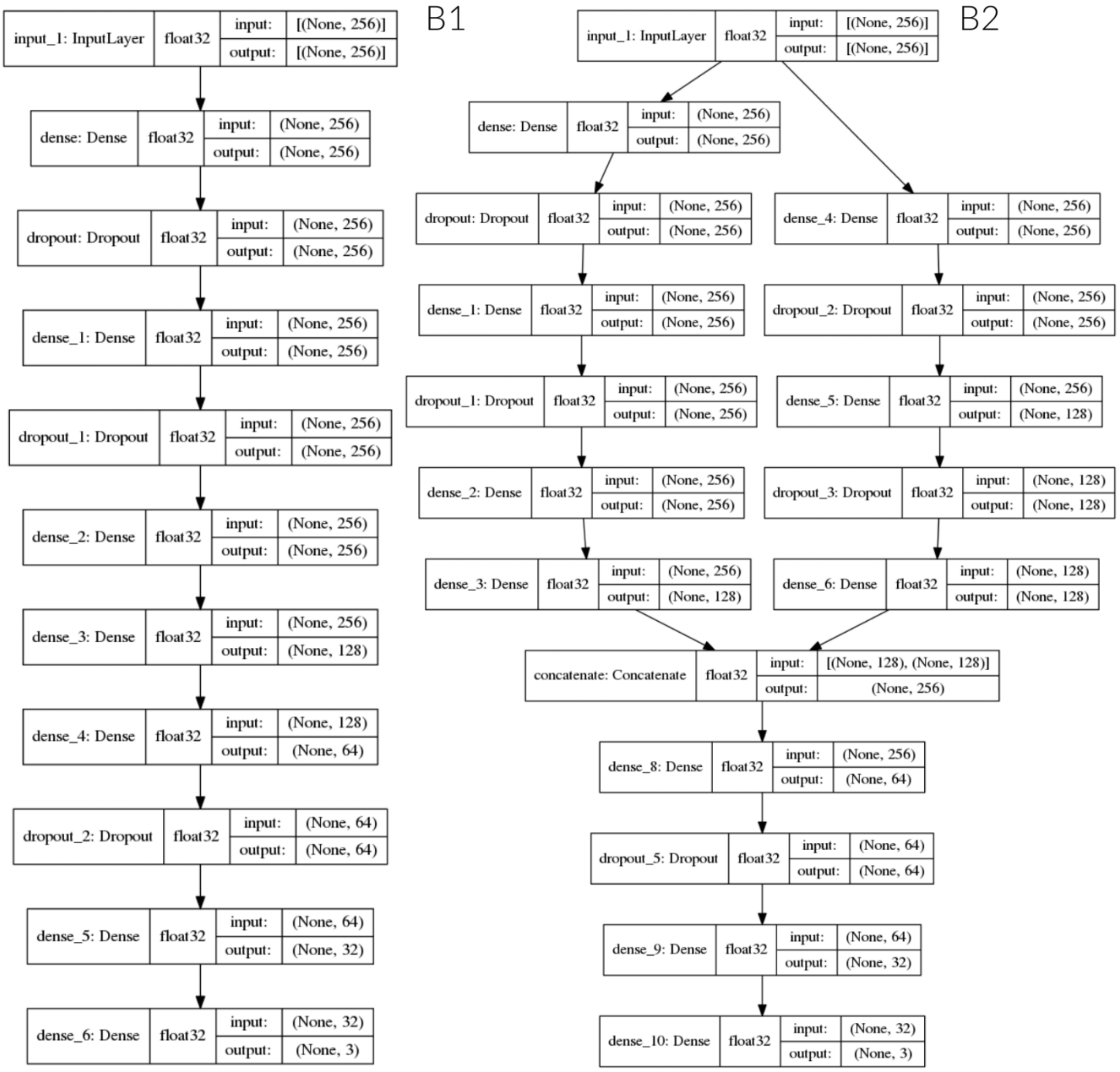


**Fig. S1.** Graph representation of the neural networks B1 and B2. Each layer is represented by a wide rectangle showing the layer type, the data type of the neurons (which is float32 for all layers), as well as the input and output shape of that layer. The output shape corresponds to the number of neurons in the layer.

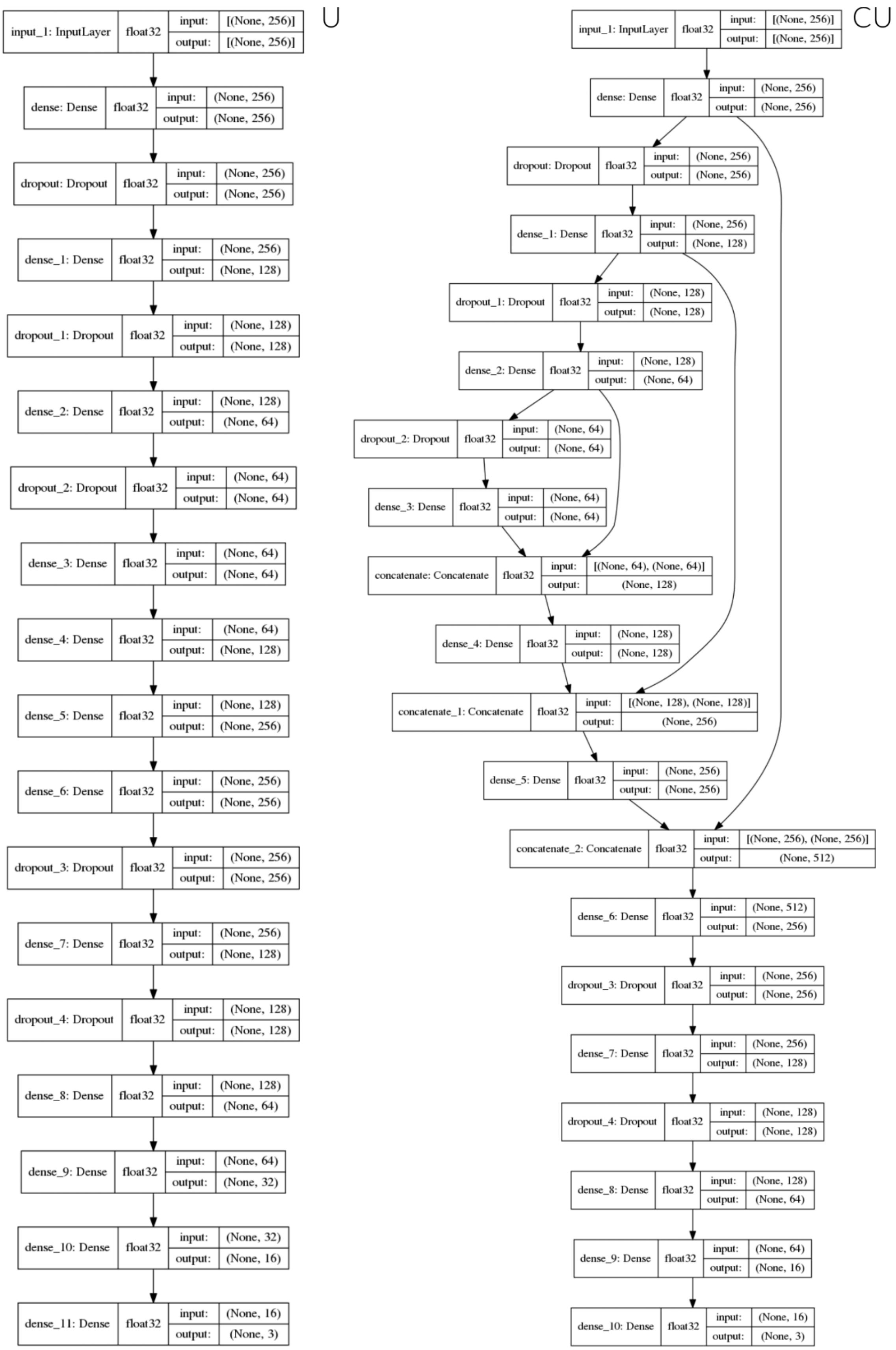


**Fig. S2.** Graph representation of the neural networks U and CU. Each layer is represented by a wide rectangle showing the layer type, the data type of the neurons (which is float32 for all layers), as well as the input and output shape of that layer. The output shape corresponds to the number of neurons in the layer.

#### 1.5 Comparing run times of neural network classifier with ML and BioNJ

Computing the likelihood function for a given phylogenetic tree, substitution model and alignment is computationally intensive. Conducting a prediction with a trained neural network of the size we use here should require comparatively fewer operations. Therefore we conjecture that predictions with the neural network could potentially be faster than predictions with the Maximum Likelihood (ML) method. In this project we compared the computing times of our neural networks for determining the best model of sequence evolution with ModelFinder (Kalyaanamoorthy et al., 2017) as implemented in IQ-Tree (Minh, 2020), and the computing times of NNs to determine the best tree topology with the Neighbour Joining (BioNJ; Gascuel, 1997) and the ML methods as implemented in IQ-Tree version 2.0.3.

Since the Python programming language has become the standard language for machine learning applications in research, we provide our code and programs in Python, knowing that an implementation in C/C++ could be significantly faster. This disadvantage is mainly confined to loading and handling the data in our code and in the Keras library, since the Numpy and the Tensorflow libraries are mainly implemented in C++. Since counting site pattern frequencies is particularly slow in Python we developed two alternative approaches:

1. Computing site pattern frequencies by calling the external quartet-pattern-counter-v1.1 program, implemented in C++.
2. Implementing the quartet pattern counter in a C module using the pybind11 library (Wenzel, 2017) such that the pattern counter can be called from within the python program. This avoids the overhead of calling an external program. It is a bit faster for nucleotide alignments and is a bit slower for amino acid alignments.

Computation time comparisons have been orchestrated by a single Python program. All computations have been done on a High Performance Computing cluster without GPU in a single run so that all computations are carried out at approximately the same time on the same CPU. Both the neural network and IQ-Tree have specifically been given a single core for their computations, in order to make the results comparable. Using a GPU should result in a significant speed improvement for the machine learning based classifier over the results we present here.

Neural networks have large file sizes and loading them requires a considerable time. Our computing times for neural networks do not include the time needed to load the neural network into the RAM, since we assume that multiple predictions are typically carried out in one program run.

**Computing times for substitution model predictions:**

We simulated 100 alignments of length 1,000, 10,000, 100,000 on a random tree topology for each nucleotide (JC+I+G, K2P+I+G, F81+I+G, HKY+I+G, GTR+I+G) and amino acid (DAY+I+G, JTT+I+G, WAG+I+G, LG+I+G) substitution model with PolyMoSim, version 1.1.4. These are in total 500 nucleotides and 400 amino acid alignments for each alignment length. From within a Python program the best substitution models were predicted and the computing times were determined for our neural networks and for ModelFinder as implemented in IQ-Tree version 2.0.3. When using the neural networks, all predictions have been done in independent calls to the predict function. Combining multiple predictions into one single call to the predict function could speed up the neural network prediction tremendously but would not provide a fair comparison with other programs. In all cases IQ-Tree determines parameters that our neural networks do not determine, such as branch lengths and model parameters. In order to reduce the resulting disadvantage when determining computation times we restrict the models IQ-Tree compares as much as possible. For determining the best nucleotide model of sequence evolution we specify the following IQ-Tree parameters: `-T 1 -m MF -mset JC+I+G,K2P+I+G,F81+I+G,HKY+I+G,GTR+I+G -redo` and for the best amino acid model of sequence evolution `-m MF -T 1 -mset Dayhoff+I+G,JTT+I+G,WAG+I+G,LG+I+G -redo`.

**Computing times for topology predictions:**

Since we introduced neural networks for determining the best tree topology separately for each model of sequence evolution, the prediction times were also determined for each substitution model. We simulated 100 alignments of lengths 1,000, 10,000, 100,000 for each substitution model and for each of the three tree topologies. These are 300 alignments for each alignment length. Now, for each substitution model we determined the best tree topology with our neural networks, with BioNJ as implemented in IQ-Tree version 2.0.3 using the options `-fast -T 1 -m Correct-Model -redo` and with the ML method implemented in IQ-Tree with the options `-T 1 -m Correct-Model -redo`, where Correct-Model was one out of {JC, F81, F84, K2P, HKY, GTR} for nucleotide or {Dayhoff, JTT, WAG, LG} for amino acid data sets.

Even though we determine computing times separately for each of the six neural network architectures, in Figure S3 we only provide the computing time for the NN that had the highest accuracy for the corresponding prediction task.

#### 1.6 Utility of alternative machine learning classifiers including support vector machine and random forest classifiers

Simple frequency distributions are often classified with non-neural network classifiers such as the naive gaussian, support vector machine, or random forest classifiers (Hastie et al., 2009). Potentially, these classifiers could be suited equally well or even better than neural network classifiers to predict the best tree topology from site pattern frequencies. Their main advantage is that they are less computationally demanding than neural networks. Even though our main focus lies on exploring the utility of neural networks we also want to test the general utility of these classifiers for topology predictions.

We employed the lazypredict library version 0.2.12 (Pandala, 2020) which automatically tests and compares a large set of non-neural-network classifiers implemented in the Scikit-learn library (Pedregosa et al., 2011). A full list of the classifiers included in this comparison is given in Figure S4. To train these classifiers and to assess their accuracy in predicting the phylogenetic tree topologies for nucleotide alignments, 600,000 alignments of length 100,000 bp were simulated. Sequences were allowed to evolve under the K2P+I+G substitution model under all three topologies with branch lengths and model parameters chosen as for the training data of the neural networks (see Methods in main text). Half of the alignments were simulated on trees with short ($0.001<v<0.02$) and the other half on trees with normal ($0.1<v<0.5$) internal branch lengths. For amino acid alignments, training and accuracy of the topology prediction were assessed using 6,000 simulated alignments of length 1,000,000 aa that evolved under the Dayhoff+I+G substitution model under all three topologies. Branch lengths and model parameters were chosen as before. Training was performed on 96% of these alignments, while the validation was conducted on the remaining 4%. When using the lazypredict library, one is restricted to the same alignment length in the training and validation data set. For a comparison of these results with the neural network topology prediction success, we created simulated verification alignments of the same lengths and created with the same characteristics as above. The Python program that conducts the comparison of machine learning classifiers with the lazy predict library is available in the DeepNNPhylogeny Github repository under the link DeepNNPhylogeny/-lazypredict.

### 2. Supplementary results and figures

#### 2.1 Run time of machine learning classifier

Computation time results are illustrated in Figure S3. They show that the neural network classifiers are much faster than ModelFinder in determining the best model of sequence evolution and also much faster than IQ-Tree in determining the best ML tree topology. The ML method consistently requires computing times for model and topology predictions that are at least a factor of 3-4 larger than neural networks for nucleotide alignments and a factor of 16 larger for amino acid alignments of length 1,000. The advantage for neural networks is larger for amino acid alignments than for nucleotide alignments and it gets larger the longer the alignments are. For amino acid alignments of length 100,000, neural networks are 1300 times faster for determining the best model of sequence evolution and 247 times faster for determining the best tree topology. Neural networks are also faster than the BioNJ method in determining the best tree topology, except for nucleotide alignments of length 1,000 bp.

It should be noted that for determining the best ML tree and the best BioNJ tree, IQ-Tree computes parameters such as branch lengths and model parameters that are not determined by our neural networks.

From repeating the analyses multiple times we see an approximate variation of the computing times that is well below ±10% from one run to the next. We refrained from quantifying the exact standard deviation, since it would require considerable computational resources to determine it.

Figure S3 visualises the prediction times for that neural network architecture which showed the highest prediction accuracy. At this point it should be noted that the prediction times differ only marginally among the different neural network architectures. Interestingly, even the prediction times for the most complex models B10 and CU are not necessarily longer than those of the other architectures.

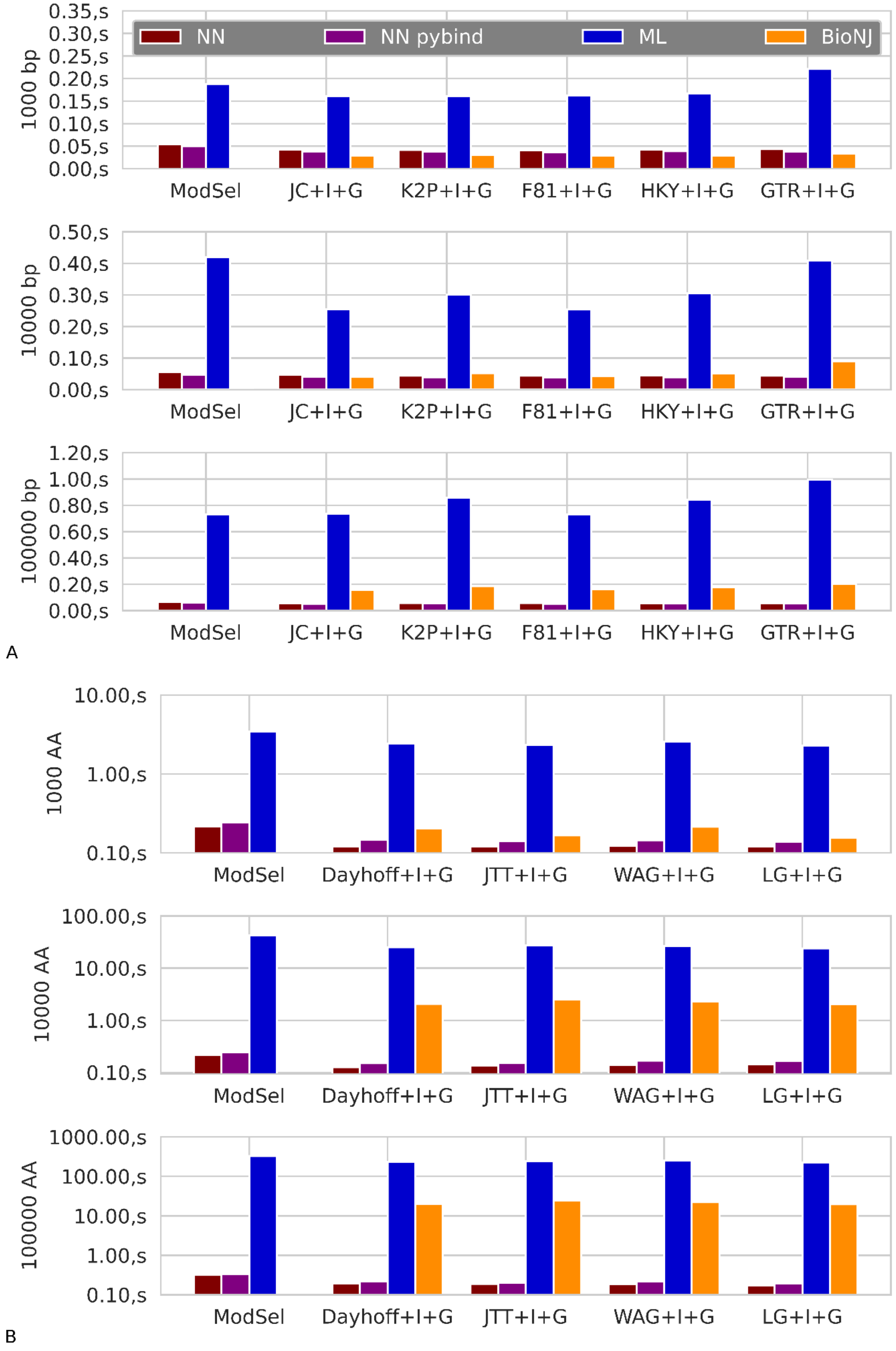

NN
NN pybind
ML
BioNJ
1000 bp
10000 bp
100000 bp
0.35,s
0.30,s
0.25,s
0.20,s
0.15,s
0.10,s
0.05,s
0.00,s
0.50,s
0.40,s
0.30,s
0.20,s
0.10,s
0.00,s
1.20,s
1.00,s
0.80,s
0.60,s
0.40,s
0.20,s
0.00,s
ModSel
JC+I+G
K2P+I+G
F81+I+G
HKY+I+G
GTR+I+G
A
1000 AA
10000 AA
100000 AA
1000.00,s
100.00,s
10.00,s
1.00,s
0.10,s
ModSel
Dayhoff+I+G
JTT+I+G
WAG+I+G
LG+I+G
B

**Fig. S3:** Mean prediction time of NNs and IQ-Tree for the model selection (ModSel) and topology prediction (model+I+G) tasks. (A) The test alignments are 1,000 bp, 10,000 bp and 100,000 bp long**.** (B) The test alignments are 1,000 aa, 10,000 aa, 100,000 aa long. A logarithmic scaling is used on the Y-axis.

#### 2.2 Classification success of alternative machine learning algorithms

Prediction accuracies for the best tree topology determined with non-neural network classifiers are illustrated in Figure S4 for nucleotide and amino acid alignments. The best performing classifier for nucleotide data is the support vector machine classifier (SVC≙SVM) with a prediction accuracy of 96%. For sequences simulated in the same way as those usFed for training and testing the lazypredict classifiers, the neural network classifier achieved an accuracy of 95.4% (B2 model). For amino acid alignments the LGBM classifier achieved the highest accuracy of 98% and the best neural network achieved an accuracy of 99.1% (B10).

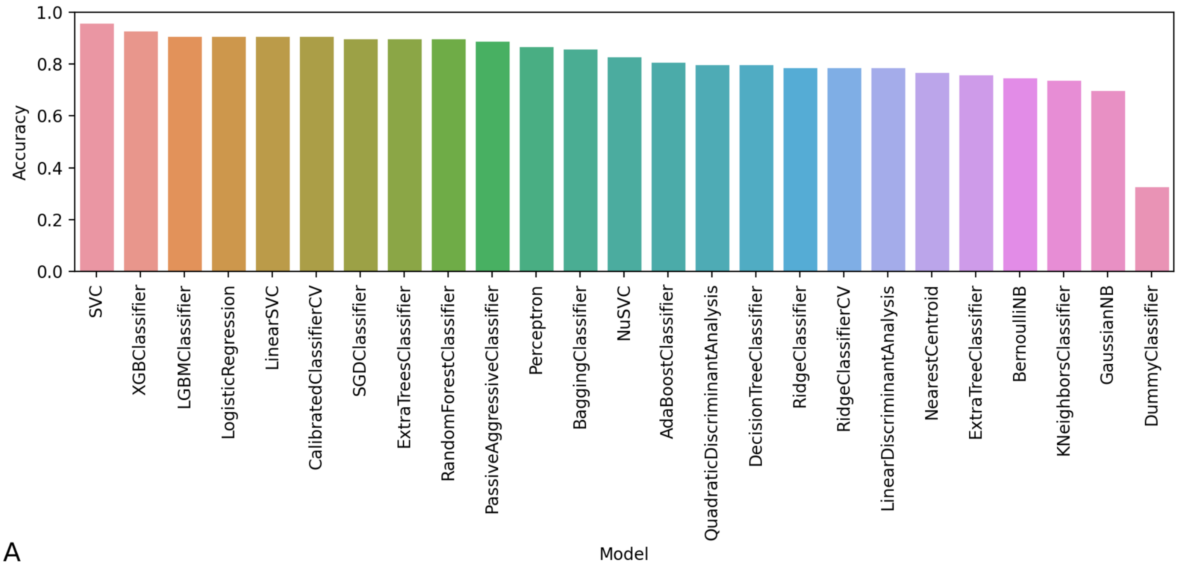


A

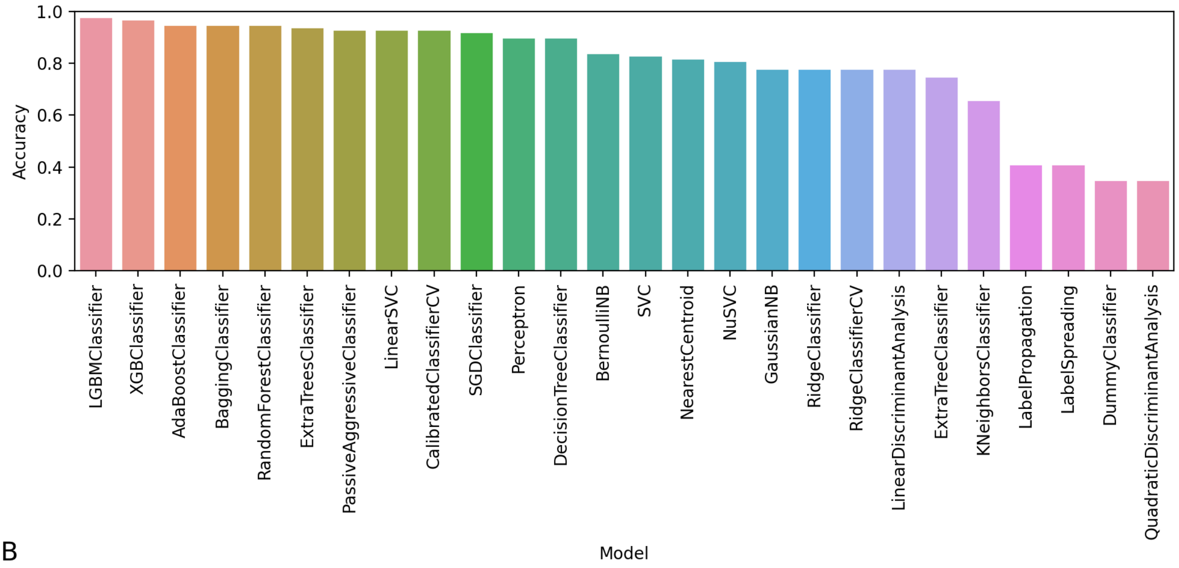


B

**Fig. S4.** The accuracy of topology predictions utilising machine learning algorithms implemented via the python library lazypredict. (A) Prediction results for 25 non-neural-network machine learning classifiers trained on a large set of nucleotide alignments. (B) Prediction results for 26 machine learning classifiers trained on a large set of amino acid alignments.

We observe that for nucleotide data with a small number of features the support vector machine classifiers (Noble, 2006) perform best, while for amino acid data with a large number of features the Light Gradient Boosting Machine classifier (LGBMClassifier; Ke et al., 2017) yield the best results.

#### 2.3 Detailed classification accuracies of the neural network classifiers compared to the ML and BioNJ method

We have illustrated the model and topology prediction accuracies of neural networks in comparison to the ML and the BioNJ methods in the main text in Figs. 3-7. Detailed prediction accuracies are given in Table S1. In particular this table lists the best performing neural network architectures together with the neural network architectures that are not significantly worse compared to the best architecture. Statistical significance was determined with a binomial test that used the standard Wilson score interval with continuity correction (Wilson, 1927; Crawley, 2014).

Table.S1. Detailed list of model and topology prediction accuracies for normal (nb) and short (sb) internal branch lengths. Bold and italic text in column "p-value ML" depicts accuracies that are significantly different. **Bold:** NN is significantly better or equally good compared to the ML method. *Italic* - NN is significantly worse than the ML method. The first two lines list model prediction accuracies and the remaining lines list topology prediction accuracies. It should be noted that a Bonferroni correction of the level of significance would remove several of these significant results.

| **Model, Alignment length, Internal branch length** | **Best NN** | **Best NN prediction success [%]** | **ML prediction success [%]** | **BioNJ prediction success [%]** | **p-value ML** | **p-value BioNJ** | **List of not significantly worse NNs** |
|---|---|---|---|---|---|---|---|
| **Nucleotide substitution model prediction, 30000, nb** | B10 | 92.6 | 87.1 | | **<2e-16** | - | B1, B2, B3, CU |
| **Amino acid substitution model prediction accuracies. Alignment length: 10000, nb** | B10 | 99.8 | 100 | | *0.01* | - | B1, B2, B3, U |
| **JC, 1000, nb** | B10 | 99.5 | 99.5 | 72.5 | **0.86** | **<2e-16** | all |
| **K2P, 1000, nb** | B1&B3 | 99.3 | 99.4 | 87.8 | **0.63** | **<2e-16** | all |
| **F81, 1000, nb** | B1 | 99.6 | 99.6 | 73.6 | **1.00** | **<2e-16** | U, CU |
| **F84, 1000, nb** | CU | 99.6 | 99.5 | 87.3 | **0.71** | **<2e-16** | all |
| **HKY, 1000, nb** | CU | 99.5 | 99.5 | 88.3 | **1.00** | **<2e-16** | all |
| **GTR, 1000, nb** | CU | 99.4 | 99.5 | 79.9 | **0.86** | **<2e-16** | all |
| **JC, 10000, nb** | B1,B2,B3 | 100 | 100 | 72.9 | **1.00** | **<2e-16** | all |
| **K2P, 10000, nb** | B3&CU | 100 | 100 | 87.0 | **1.00** | **<2e-16** | all |
| **F81, 10000, nb** | B2,B10, CU | 100 | 100 | 73.3 | **1.00** | **<2e-16** | B1,B3 |

| | | | | | | | |
|---|---|---|---|---|---|---|---|
| **F84, 10000, nb** | B1,B3, B10,CU | 100 | 100 | 86.8 | **1.00** | **<2e-16** | all |
| **HKY, 10000, nb** | B2,B3, CU | 99.97 | 100 | 86.8 | **1.00** | **<2e-16** | B1,B10 |
| **GTR, 10000, nb** | B1 | 100 | 100 | 81.8 | **1.00** | **<2e-16** | all |
| **JC, 1000, sb** | CU | 47.6 | 48.0 | 38.4 | **0.76** | **1e-12** | all |
| **K2P, 1000, sb** | B3 | 48.3 | 48.0 | 42.7 | **0.84** | **2e-05** | all |
| **F81, 1000, sb** | B10 | 48.2 | 47.8 | 39.2 | **0.74** | **3e-12** | all |
| **F84, 1000, sb** | B3 | 47.9 | 48.3 | 43.3 | **0.77** | **3e-4** | all |
| **HKY, 1000, sb** | B1 | 48.1 | 47.9 | 43.3 | **0.94** | **2e-4** | all |
| **GTR, 1000, sb** | B3 | 47.9 | 48.4 | 43.3 | **0.76** | **3e-4** | all |
| **JC, 10000, sb** | B3 | 71.2 | 73.4 | 40.4 | **0.06** | **<2e-16** | B1,B2,B10,CU |
| **K2P, 10000, sb** | B2&B10 | 70.2 | 72.9 | 49.0 | *0.03* | **<2e-16** | B1,B3,CU |
| **F81, 10000, sb** | B2 | 70.1 | 73.3 | 40.1 | *0.01* | **<2e-16** | B1,B3,B10,CU |
| **F84, 10000, sb** | B2 | 71.3 | 73.6 | 49.3 | *0.04* | **<2e-16** | B1,B3,B10,CU |
| **HKY, 10000, sb** | B2 | 71.0 | 73.2 | 49.2 | **0.07** | **<2e-16** | B1,B3,B10,CU |
| **GTR, 10000, sb** | B2 | 70.6 | 71.8 | 43.2 | **0.31** | **<2e-16** | B1,B3,B10 |
| **JTT, 1000, nb** | B3 | 98.2 | 99.9 | 92.6 | *1e-11* | **<2e-16** | B2, U |
| **LG, 1000, nb** | B3 | 98.3 | 99.9 | 93.0 | *2e-10* | **<2e-16** | U |
| **WAG, 1000, nb** | B3 | 98.2 | 99.97 | 91.6 | *3e-12* | **<2e-16** | B2 |
| **DAY, 1000, nb** | B3 | 97.5 | 99.9 | 92.5 | *1e-15* | **<2e-16** | B2, U |
| **JTT, 1000, sb** | B3 | 43.5 | 60.2 | 46.2 | *<2e-16* | *0.036* | B2,U,CU |
| **LG, 1000, sb** | B3 | 41.2 | 60.2 | 47.4 | *<2e-16* | *3e-06* | B1,B2,U,CU |
| **WAG, 10000, sb** | B3 | 43.2 | 60.1 | 46.7 | *<2e-16* | *0.007* | B2,CU |
| **DAY, 1000, sb** | B3 | 43.1 | 59.9 | 47.3 | *<2e-16* | *0.001* | B2,U,CU |
| **JTT, 10000, nb** | B3&CU | 99.9 | 100 | 93.6 | **0.25** | **<2e-16** | B1,B2,U |
| **LG, 10000, nb** | B3 | 99.9 | 100 | 94.7 | **0.48** | **<2e-16** | B1,B2,CU |
| **WAG, 10000, nb** | B3 | 99.8 | 100 | 93.1 | **0.07** | **<2e-16** | B1,B2,U,CU |
| **DAY, 10000, nb** | B3 | 99.8 | 100 | 93.8 | *0.023* | **<2e-16** | B1,B2,U,CU |
| **JTT, 10000, sb** | B3 | 59.5 | 87.7 | 61.2 | *<2e-16* | 0.205 | none |
| **LG, 10000, sb** | B3 | 58.5 | 87.3 | 62.1 | *<2e-16* | *0.006* | B2,CU |
| **WAG, 10000, sb** | B3 | 60.2 | 87.0 | 60.0 | *<2e-16* | 0.874 | none |
| **DAY, 10000, sb** | B3 | 58.1 | 86.6 | 60.6 | *<2e-16* | *0.046* | CU |
